# Free-energy-based framework for early forecasting of stem cell differentiation

**DOI:** 10.1101/692285

**Authors:** H. Suresh, S.S. Shishvan, A. Vigliotti, V.S. Deshpande

## Abstract

Commitment of stem cells to different lineages is inherently stochastic but regulated by a range of environmental bio/chemo/mechanical cues. Here we develop an integrated stochastic modelling framework for predicting the differentiation of hMSCs in response to a range of environmental cues including sizes of adhesive islands, stiffness of substrates and treatment with ROCK inhibitors in both growth and mixed media. The statistical framework analyses the fluctuations of cell morphologies over around a 24-hour period after seeding the cells in the specific environment and uses the distribution of their cytoskeletal free-energy to forecast the lineage the hMSCs will commit to. The cytoskeletal free-energy which succinctly parameterises the biochemical state of the cell is shown to capture hMSC commitment over a range of environments while simple morphological factors such as cell shape, tractions on their own are unable to correlate with lineages hMSCs adopt.

## Introduction

Stem cells have the dual ability to differentiate into various mature cells (such as osteoblasts, chondrocytes, neuroblasts, etc.) that form various tissues, and proliferate to maintain a pool of immature cells that can differentiate when required. The degree and outcome of differentiation are controlled by various extrinsic signals in the stem cell niche that include cell-cell, cell-matrix and cell-soluble cue interactions [1, 2]. For example, human mesenchymal stem cells (hMSCs) choose adipogenic over osteogenic lineage when plated at a high cell density, where the extent of cell-cell interactions determines cell fate [3]. Creation of synthetic cellular niches requires careful choices for the extracellular matrix (ECM), and soluble factors (such as growth factors, cytokines and hormones) to best harness the regenerative potential of stem cells [2, 4].

While the effect of soluble factors on stem cell lineage commitment and differentiation has been extensively studied, a thorough investigation of the influence of insoluble signals such as extracellular matrix rigidity, and adhesive properties of the substrate is still ongoing. It is now well-known that environmental cues such as microgravity [5] and mechanical cues such as substrate rigidity, substrate curvature, arrangements of micropillars, gratings and wells [6-9] dictate cell fate. Nanoscale physical cues such as nanotubes and nanowires of different pore sizes and spacing, nano-grating, nano-posts, and different arrangements of nano-pits [10-14] act at the scale of single focal adhesions to set cell lineage. Advances in nano- and micropatterning have aided in exploring the effect of chemical cues (such as changing the concentration and spacing of adhesive proteins on the substrate [15]) and geometric cues (i.e., confining the cells to adhesive patterns of different shapes and sizes [3, 16]) on cell fate. Over the past two decades, numerous experiments have been performed to investigate the effects of mechano/chemo/geometric cues (and their combinations) on the differentiation of multipotent stem cells. At the same time, several models have also been developed and refined using the wealth of information provided by experiments.

One class of models simulate cell shape, cytoskeletal arrangements and focal adhesion formation in cells in response to the physical cues in the ECM, and thereby predict cell differentiation. Qualitative predictions of cell shape-induced differentiation on elastic substrates are obtained using 3D finite element models, where relevant subcellular structures are modelled explicitly [17]. Cell-cell interactions are captured through discrete finite element models (where each cell is a discrete unit that interacts with other cells and the substrate through contact stresses), and the extent of cell deformation is correlated with degree and lineage of differentiation [18]. Spreading of cells on patterned substrates are modelled using particle-based methods, with inter-particle forces chosen to best capture the pinning of cells on substrates by the adhesive islands [19]. While these coarse-grained models provide an intuitive picture of the mechanisms of cell spreading and differentiation, they are often tailored for specific cues.

Another class of models use machine-learning techniques to detect patterns in large volumes of experimental observations that can enable cell fate prediction. The experimental observations can be related to the expression of transcription factors (such as CBF*α*1 for osteoblasts and PPAR*γ* for adipocytes) [20, 21], or various measures of cell shape (such as cell area, aspect ratio, stress-fibre intensity, etc.) [22, 23]. However, a frequent output from these models is a complex combination of input variables that seems to correlate strongly with cell fate, but physical significance of such measures remain poorly understood. Ideas borrowed from statistical mechanics have also been widely used to relate inherently stochastic molecular fluctuations to well-defined macroscopic cell fates [24, 25], but such models do not provide insight into cell shape changes associated with the different stages of the differentiation process.

Here we aim to combine the strengths of the different approaches into one integrated stochastic framework for stem cell differentiation. While we recognise that cells exist in a fluctuating equilibrium with their surroundings, we also realise the need to incorporate fluctuations to the cell shape and cytoskeletal structure (rather than at the gene-level) so that we can predict both the spreading and differentiation response of hMSCs to chemo-mechanical cues in the ECM. In this study, we present a framework to (i) predict cell differentiation over mechanical and chemical cues, and (ii) show the equivalence of different types of chemo-mechanical cues in directing cell lineage commitment and subsequent differentiation.

## Modelling

While it is well-established that changes to gene expressions over a period of about 1-2 weeks dictate the lineage commitment and differentiation fate of hMSCs, more recent studies have indicated that a combination of a large number of cell, nuclear and cytoskeletal morphometrics also provides excellent forecasting of the lineage of hMSCs [22, 23]. These morphometrics develop over a period of 1 to 2 days when the gene expression of cells has not been affected irreversibly by the environment. Here, we apply the recently developed *homeostatic mechanics* to predict the distribution of morphological states the cell assumes in the interphase period of its cell cycle, which in turn relates to its differentiation outcome. The homeostatic mechanics framework has already been shown to successfully capture a range of observations for smooth muscle cells seeded on elastic substrates [26, 27] and for myofibroblasts seeded on substrates micropatterned with stripes of fibronectin [28], giving us confidence to investigate its generality in terms of predicting the differentiation of hMSCs in response to a range of environmental cues.

### 2.1 A brief overview of the homeostatic mechanics framework

The homeostatic mechanics framework recognises that the cell is an open system which exchanges nutrients with the surrounding nutrient bath (Fig. 1a). These high-energy nutrient exchanges fuel large fluctuations (much larger than thermal fluctuations) in cell response associated with various intracellular biochemical processes. However, these biochemical processes attempt to maintain the cell in a homeostatic state, i.e. the cell actively maintains itself out of thermodynamic equilibrium [29] by maintaining its various molecular species at a specific average number over these fluctuations that is independent of the environment [30]. More specifically, homeostasis is the ability of a living cell to maintain, via coupled and inter-connected biomechanical processes, the concentration of all internal species at a fixed average value independent of the environment, over all its non-thermal fluctuations. Upon employing the homeostatic constraint, we have the result that the ensemble average Gibbs free-energy is equal to that of the cell in suspension. This is a universal constraint that quantifies the fact that living cells maintain themselves away from thermodynamic equilibrium but yet attain a stationary state. Recognising that biochemical processes such as actin polymerisation and treadmilling provide the mechanisms to explore morphological microstates, we employ the *ansatz* that the observed distribution of cell shapes is that one with the overwhelming number of microstates, i.e. the distribution that maximises the *morphological entropy* subject to the homeostatic constraint and any other geometrical constraints such as confinement imposed by patterning adhesive islands on substrates.

**Figure 1:**
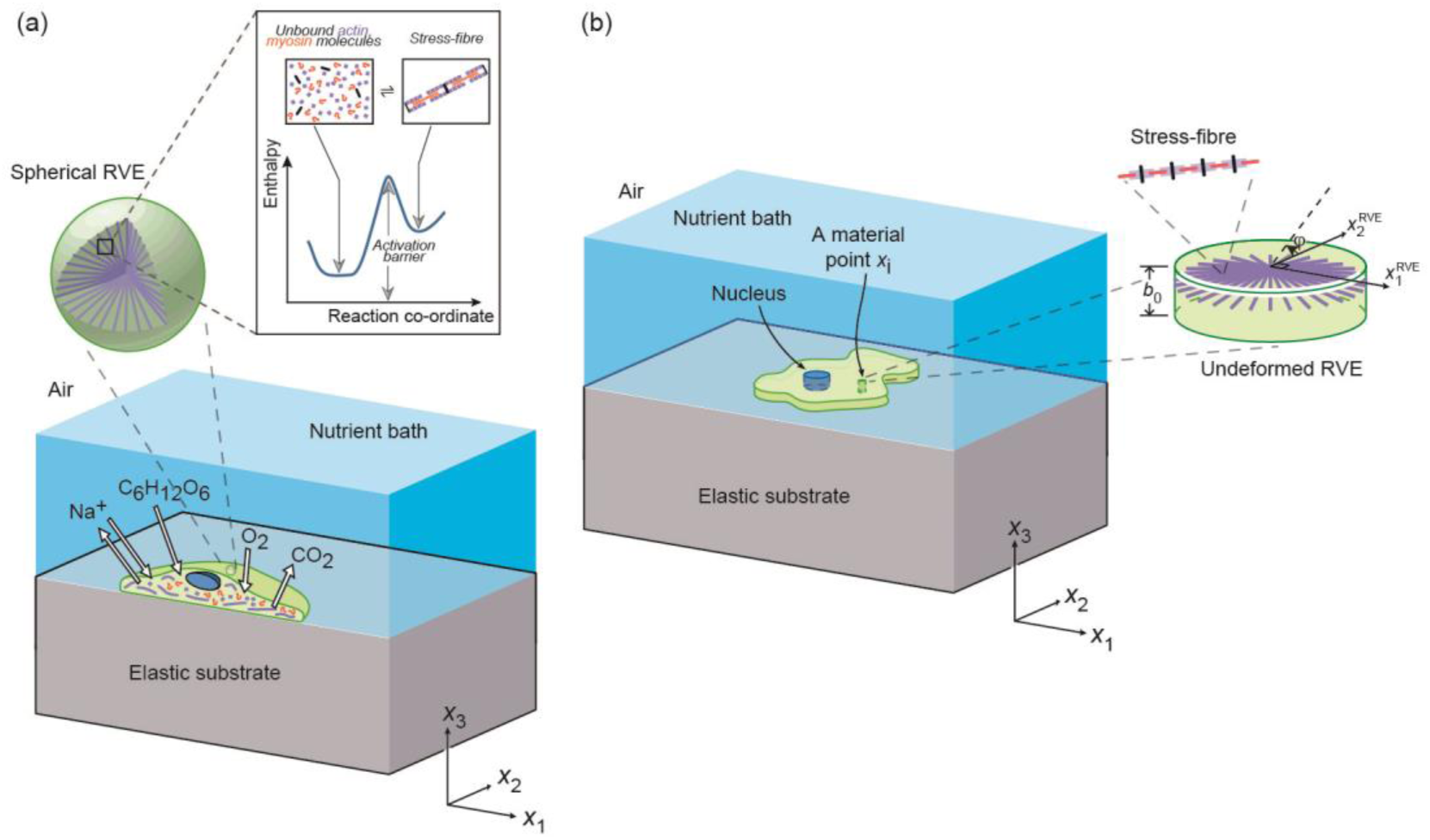
(a) An illustration of the cell model employed in the simulations using the homeostatic mechanics framework. The sketch shows a section of a cell on an elastic substrate and exchanging species with the nutrient bath. The inset shows a representative volume element (RVE) of the cell cytoplasm containing polymerised acto-myosin stress-fibres and the unbound proteins along with the energy landscape that governs the equilibrium of these proteins. (b) The two-dimensional (2D) approximation of the cell with the 2D RVE.

Shishvan et al. [26] obtained the equilibrium distribution of states that the cell assumes in terms of the Gibbs free-energy *G*^(*j*)^ of the morphological state (*j*) of the system as

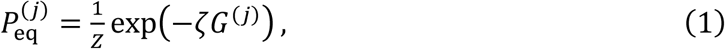

where *Z* ≡ Σ_*j*_exp(−*ζ G*^(*j*)^) is the partition function of the morphological microstates, and the distribution parameter *ζ* follows from the homeostatic constraint 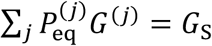, where *G*_s_ is the equilibrium Gibbs free-energy of an isolated cell in suspension. Thus, 1/*ζ* in (1) is referred to as the *homeostatic temperature* that is conjugated to the morphological entropy of the cell. We employ Markov Chain Monte Carlo to construct a Markov chain that is representative of the homeostatic ensemble. This involves calculation of *G*^(*j*)^ for a given morphological microstate (*j*) and construction of a Markov chain that is representative of the ensemble of states with probability distribution (1). Typical Markov chains comprised in excess of 2 million spread states (a detailed overview of the procedure is provided in Supplementary S1.2).

### 2.2 Gibbs free-energy of a morphological microstate

The implementation of the homeostatic mechanics approach described above requires a specific model for the Gibbs free-energy of the cell-substrate system in a given morphological state. Modelling all the elements of the cell is unrealistic and often not required as specific components are known to strongly respond to different cues. Here we are interested in investigating the differentiation behaviour of hMSCs to mechanical cues provided by the substrate stiffness [6] and geometric cues imposed by the size of adhesive islands patterned on substrates [3]. These cues are known to result in significant remodelling of the stress-fibre cytoskeleton and thus here we use a model for the Gibbs free-energy developed by Vigliotti et al [31] and subsequently modified in [26-28]. Details of the model including the parameters are given in Supplementary S1.3 and here we give a brief overview.

Single human mesenchymal stem cells (hMSCs) are modelled as two-dimensional bodies in the *x*_1_ − *x*_2_ plane lying on the surface of an elastic substrate such that the out-of-plane Cauchy stress *Σ*_33_ = 0 (Fig. 1b). The substrate is modelled as a linear elastic half-space, whereas the cell consists of only three components: a cytoplasm that is modelled as comprising an active stress-fibre cytoskeleton wherein the actin and myosin proteins exist either in unbound or in polymerised states (Fig. 1a), a passive elastic nucleus, and elements such as the cell membrane, intermediate filaments and microtubules that are all lumped into a single passive elastic contribution. The cell in its undeformed state is a circle of radius *R*_0_ with a circular nucleus of radius *R*_N_ whose centre coincides with that of the cell; see Supplementary S1.3 for details including the cell parameters used to characterise hMSCs.

For a given morphological microstate, the strain distribution within the cell is specified which directly gives the elastic strain energy of the cell 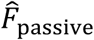 via a 2D Ogden-type hyperelastic model for both the nucleus and cytoplasm. The stress-fibre cytoskeleton within the cytoplasm is modelled as a distribution of stress-fibres such that at each location *x*_*i*_ within the cell, 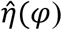 parameterises the angular concentration of stress-fibres over all angles *φ*, while 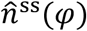 denotes the number of functional units within each stress-fibre. Thus, at any *x*_*i*_ there is a total concentration 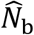 of bound stress-fibre proteins obtained by integrating 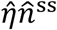 over all orientations *φ*, and these bound proteins are in chemical equilibrium with the unbound stress-fibre proteins (Fig. 1a). The unbound proteins are free to diffuse within the cell, and thus at equilibrium of a morphological microstate, the concentration 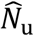 of these unbound stress-fibre proteins is spatially uniform. This chemical equilibrium condition along with the conservation of stress-fibre proteins within the cells provides the spatial and angular distributions of stress-fibres from which the free-energy of the cytoskeleton 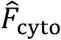 is evaluated. The tractions that the cell exerts on the substrate induces a Helmholtz free-energy 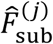 within the substrate. Then, the total (normalised) free-energy of the cell-substrate system in morphological microstate (*j*) follows as 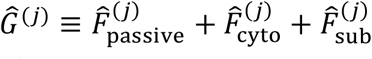 (see Supplementary S1.4 for details of the normalisations).

### 2.3 Early forecasting of the lineage of hMSCs

A combination of a large number of cell, nuclear and cytoskeletal morphometrics that develop over a period of 1-2 days have been shown to forecast the lineage of hMSCs as measured via gene expressions over a period of about 1 week [22, 23]. While such a morphometric analysis is undoubtedly useful it has two drawbacks: (i) it requires the measurement and analysis of a large number of morphological metrics, and (ii) it provides little insight into the physical phenomena that set the lineage of the cell. A number of studies have shown that the hMSCs maintain an undifferentiated state when the polymerisation of the stress-fibre cytoskeleton is inhibited by the addition of re-agents such as 2,3-butanedione monoxime (BDM) or Blebbistatin. This combined with the fact that lineage is shown to correlate with cytoskeletal morphometrics suggests that the state of the stress-fibre cytoskeleton can be used to predict cell fate. The stress-fibre cytoskeleton is modelled in detail for each morphological microstate as briefly described above and in detail in Supplementary S1.1. Of course, a number of cytoskeletal morphometrics can be extracted from the simulations much like in the experiments, but the model has the additional feature that it also quantifies the cytoskeletal free-energy 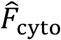 which is a succinct scalar parameter that quantifies the biochemical state of the cytoskeleton. Thus, here we attempt to use 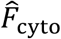 as the metric to forecast cell fate.

Unlike, deterministic free-energy models [31-33] that treat cells as systems that minimise their free-energy, the homeostatic ensemble recognises that cells exchange nutrients with their environment and thereby maintain a thermodynamic non-equilibrium but nevertheless stationary state that is commonly referred to as the homeostatic state. In this homeostatic state, hMSCs fluctuate over the equilibrium distribution of morphological microstates characterised by (1) and thus have a fluctuating 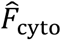 Consider the fluctuating response of a hMSC over a time period *T*_s_ when it chooses its lineage 𝕩, where 𝕩 for example could denote an osteoblast (and then subsequently goes on to differentiate to 𝕩 over a much longer time scale). Over the time *T*_s_, the average cytoskeletal free-energy of the hMSC is given by

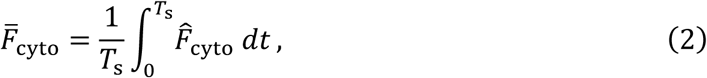

where time *t* = 0 is an arbitrary reference. We shall assume that the hMSC chooses a lineage 𝕩 if 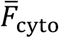 lies in the range 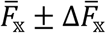, where 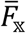 and 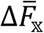 are values specific to lineage 𝕩. In order to estimate the probability of the hMSC choosing a lineage 𝕩, we first need to calculate the probability that 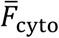 lies in the range 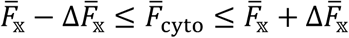. We hypothesise that cell lineage is typically set over a period of 24 to 48 hours after seeding of hMSCs in a particular environment. Over this period, the cell assumes a large number of morphological microstates. Most of these morphological microstates are correlated with each other, with the cell retaining memory of the history of its state over a timescale of 10s of minutes. However, over a longer time period, the cell loses memory and its morphological microstates are decorrelated [34]. Over the period of 24 to 48 hours when the cell chooses its lineage, we assume that hMSCs assume *N*_c_ decorrelated morphological microstates. This is thus equivalent to randomly drawing *N*_c_ morphological configurations from the homeostatic distribution (1), and then for large *N*_c_, the central limit theorem specifies that 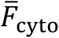 has a distribution given by the probability density function

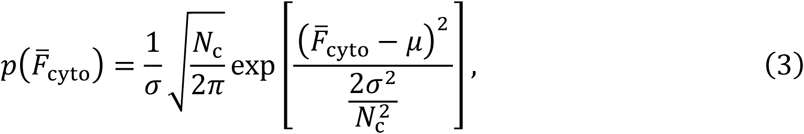

where *μ* and *σ* are the mean and standard deviations of the homeostatic distribution of 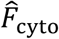, i.e. 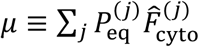 and 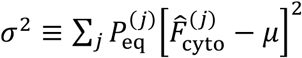 with the summations carried out over the entire homeostatic ensemble of morphological microstates (*j*). Given that a hMSC chooses a lineage 𝕩 if 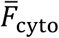 lies in the range 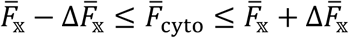, the probability *P*_𝕩_ of it choosing lineage 𝕩 is proportional to 𝒫_𝕩_ given by

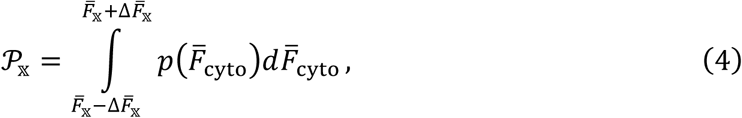

with the probability *P*_𝕩_ then defined through a normalising constant *Z*_L_ as *P*_𝕩_ ≡ 𝒫_𝕩_/*Z*_L_. This normalising constant ensures that the sum of the probabilities of all lineages with non-differentiation also treated as a lineage in the context, equals unity. Thus, *Z*_L_ ≡ max[1, Σ_𝕩_ 𝒫_𝕩_] so that if *Z*_L_ = 1 the probability of non-differentiation is vanishing and otherwise is given by 1 − Σ_𝕩_ 𝒫_𝕩_. The model for prediction of cell fate thus requires the parameter *N*_c_ in addition to 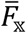 and 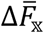 for each lineage 𝕩. Typically, these parameters are dependent also on the media in which the hMSCs are cultured and we proceed to present predictions for the differentiation of hMSCs into osteoblasts, myoblasts and adipocytes cultured in growth and mixed media. For a given media, these parameters were calibrated for a given set of cues (e.g. stiffness cues) and then used to make predictions for a different set of cues (e.g. size of adhesive islands). We emphasize that in many cues (e.g. stiffness cues), not all lineages are detected and thus calibration of the full model using experimental data is not always possible. Thus, while calibrating the model here, we lump those lineages into the “non-differentiation” category. This approximation does not add any error in the analysis so long as joint probability of differentiation of hMSCs into the detected and undetected lineages is zero. For example, consider the case of hMSCs on a range of substrate stiffness where the hMSCs can differentiate into osteoblasts, myoblasts and adipocytes but the experiment only detects osteoblasts and myoblasts. Since differentiation into adipocytes occurs only for very low stiffness values where the probability of differentiation into osteoblasts and myoblasts is vanishingly small, not accounting for differentiation into adipocytes does not add any errors for predicting the differentiation into osteoblasts and myoblasts.

## Results

We shall consider the response of hMSCs in both growth media and mixed media when seeded on elastic substrates of varying stiffness and on effectively rigid substrates patterned with adhesive islands. The response of the cells in terms of morphometrics over relatively short periods (i.e. 1-2 days) is independent of the differentiation media and thus the parameters of the hMSCs detailed in Supplementary S1.3 are not dependent on the media. However, the differentiation outcomes, and thereby 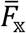 and 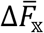 are strongly dependent on the media. We thus present results in two steps whereby we first discuss predictions of cell morphometrics and then proceed to discuss predictions of the lineage in the two different media.

### 3.1 Response on elastic substrates

The response of cells on elastic substrates is recorded through a range of observables, all of which exhibit large variations but with trends clearly emerging when the statistics of these observables are analysed. This motivates our choice of the statistical homeostatic modelling framework, in which, just as in the experimental system, observables fluctuate while trends (and understanding) emerge once these observables are viewed statistically. Figure 2a shows representative images of cell morphologies on substrates with three contrasting stiffness *E*_sub_ (while glass has a stiffness *E*_sub_ ≈ 50 GPa, the effective stiffness experienced by the cell is much lower due to the intermediate ECM, and thus following Engler et al. [6] we take *E*_sub_ = 70 kPa for glass as representing the effective stiffness of the ECM). The predictions have been presented alongside the observations (Fig. 2b) of Engler et al. [6], replicating the immunofluorescence staining used in experiments whereby the stress-fibres are shown in red, the focal adhesions in green and the nucleus in purple and blue (see Supplementary S1.4.2 for details the procedure used to translate the model predictions to such images). Overall, the cell morphologies and distributions of cytoskeletal and focal adhesion proteins are similar to the experimental observations. Stress-fibre polymerisation, focal adhesion formation, cell area and aspect ratio increase with increasing substrate stiffness, in line with a wide variety of observations [6, 8, 35]. However, as eluded to above, selected observations of cell morphologies are highly variable with understanding emerging from the statistics of observations. Recalling that typically reported observations include distributions of cell area, aspect ratio and total tractions exerted by the cell on substrates, we include in Figs. 3a-c predictions of these observables. These predictions are presented in terms of probability density functions of the normalised area *Â* (normalised by area of the cell in its elastic resting state *A*_0_), aspect ratio (via the best fit ellipse) and the normalised total tractions 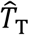 (see Supplementary S1.4.1 For details of the definitions). With increasing substrate stiffness, not only do the means of cell area, aspect ratio and total tractions increase, but so does the spread in these observables (i.e. the probability density functions are less peaked with increasing *E*_sub_).

**Figure 2:**
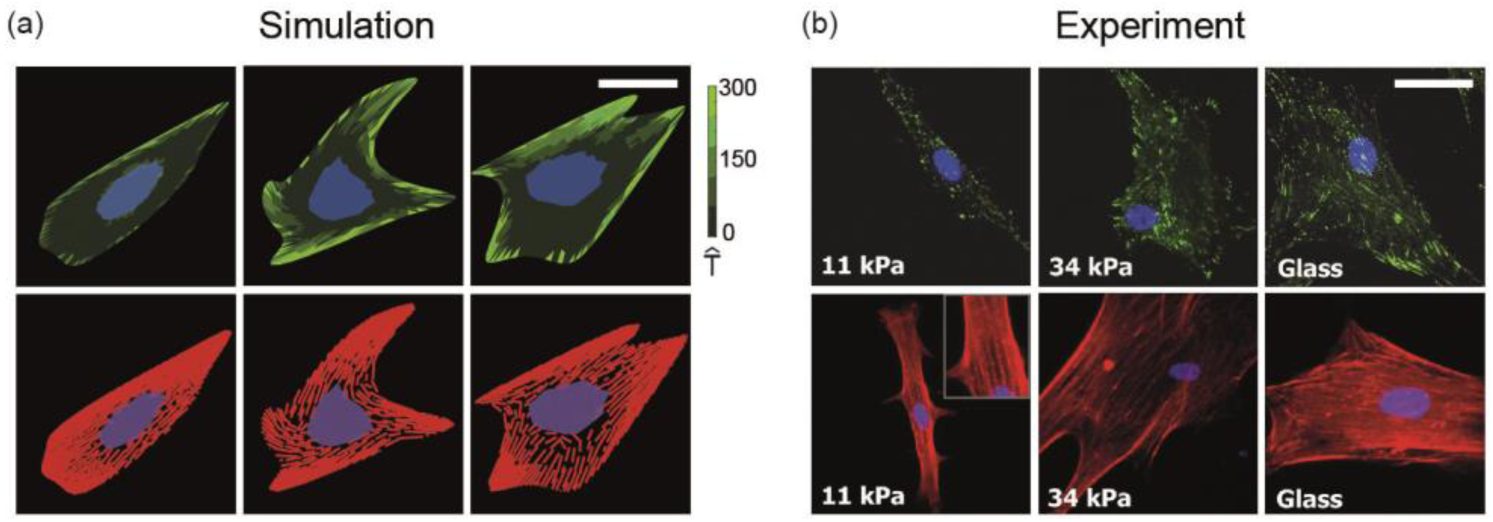
(a) Predictions from simulations, and (b) observations from Engler et al [6] of hMSCs seeded on elastic substrates uniformly coated with collagen. In the experimental immunofluorescence images, the focal adhesions are coloured green, actin red and nucleus blue and purple, and a similar scheme is followed in the predictions, with the focal adhesions parameterised by the magnitude of the normalised traction 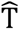 (see Supplementary S1.4.1 for details of the method used to construct immunofluorescence-like images from the simulated results). Scale bar = 20 μm.

**Figure 3:**
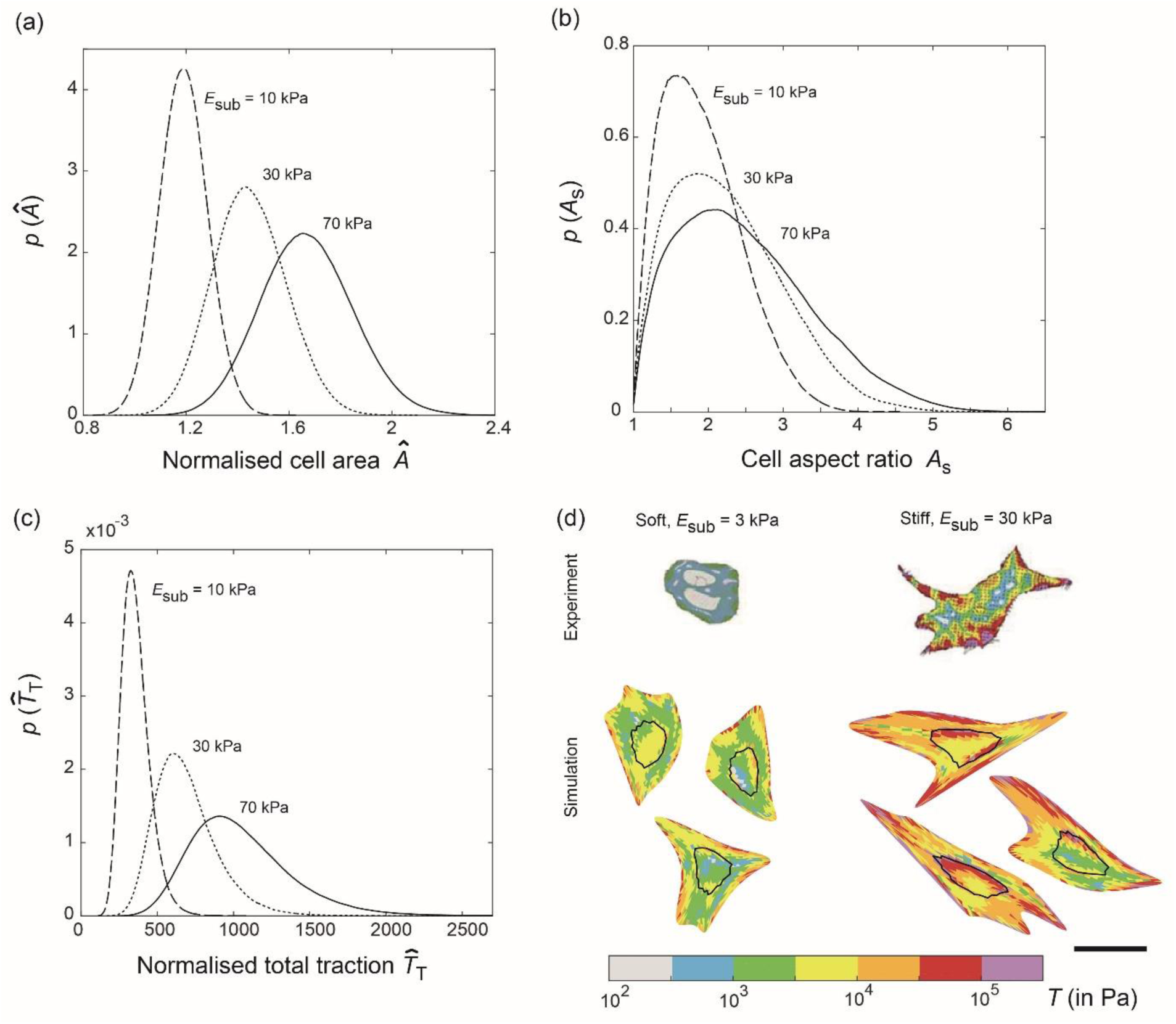
Predictions of the probability density functions of three typically reported observables for hMSCs seeded on elastic substrates uniformly coated with collagen. Distributions of (a) normalised cell area *Â*, (b) cell aspect ratio *A*_s_ and (b) normalised total traction 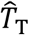 for three selected substrate stiffness *E*_sub_ in each case. (d) Comparisons between measurements [35] and predictions of the distributions of tractions T exerted by cell on elastic substrates of stiffness *E*_sub_ = 3 kPa and 30 kPa (outline of nucleus shown as a black line). In each case, we show three simulated configurations we randomly selected from the computed homeostatic distribution. The scale bar in (d) = 30 μm.

Tractions exerted by hMSCs on elastic substrates has been experimentally established, and we show in Fig. 3d predictions (for three high probability cell configurations) of the normalised traction distributions 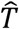 (see Supplementary S1.4.1 for definition) alongside corresponding measurements from Guvendiren & Burdick [35]. Regions of higher tractions are significantly fewer on the lower stiffness substrates, but also typically occur along the cell periphery in both the observations and predictions. In general, the agreement between model predictions of cell morphometrics and experimental observations gives confidence to attempt to employ it to aid forecasting of cell lineage. However, while substrate stiffness does affect cell response, it is clear that given the significant overlap in the distribution of observables (Figs. 3a-c), it is unlikely that these observables can be directly used to predict cell lineage.

### 3.2 Response of cell on adhesive islands

A selection of highly probable cell configurations of hMSCs on a square adhesive islands of area *A*_P_ = 2025 μm^2^ are included in Fig. 4a (these islands are patterned on PDMS substrates with *E*_sub_ = 4 MPa, which is assumed to be effectively rigid, and thus predictions shown correspond to substrates with stiffness *E*_sub_ = 70 kPa). These predictions are shown as combined immunofluorescence-like images showing stress-fibres (green), focal adhesions (pink) and nucleus (blue). Tractions exerted by hMSCs are considered to be a strong indicator of the lineage they adopt [8, 35] and we include predictions of the probability density functions of the normalised total tractions 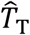 in Fig. 4b for a range of areas *A*_P_ of the square adhesive islands. Again, there is significant overlap in the traction distributions for the different island sizes, but in general, the tractions that cells exert decrease with decreasing *A*_P_, and this is generally thought to indicate a preference for differentiation into adipocytes over osteoblasts.

**Figure 4:**
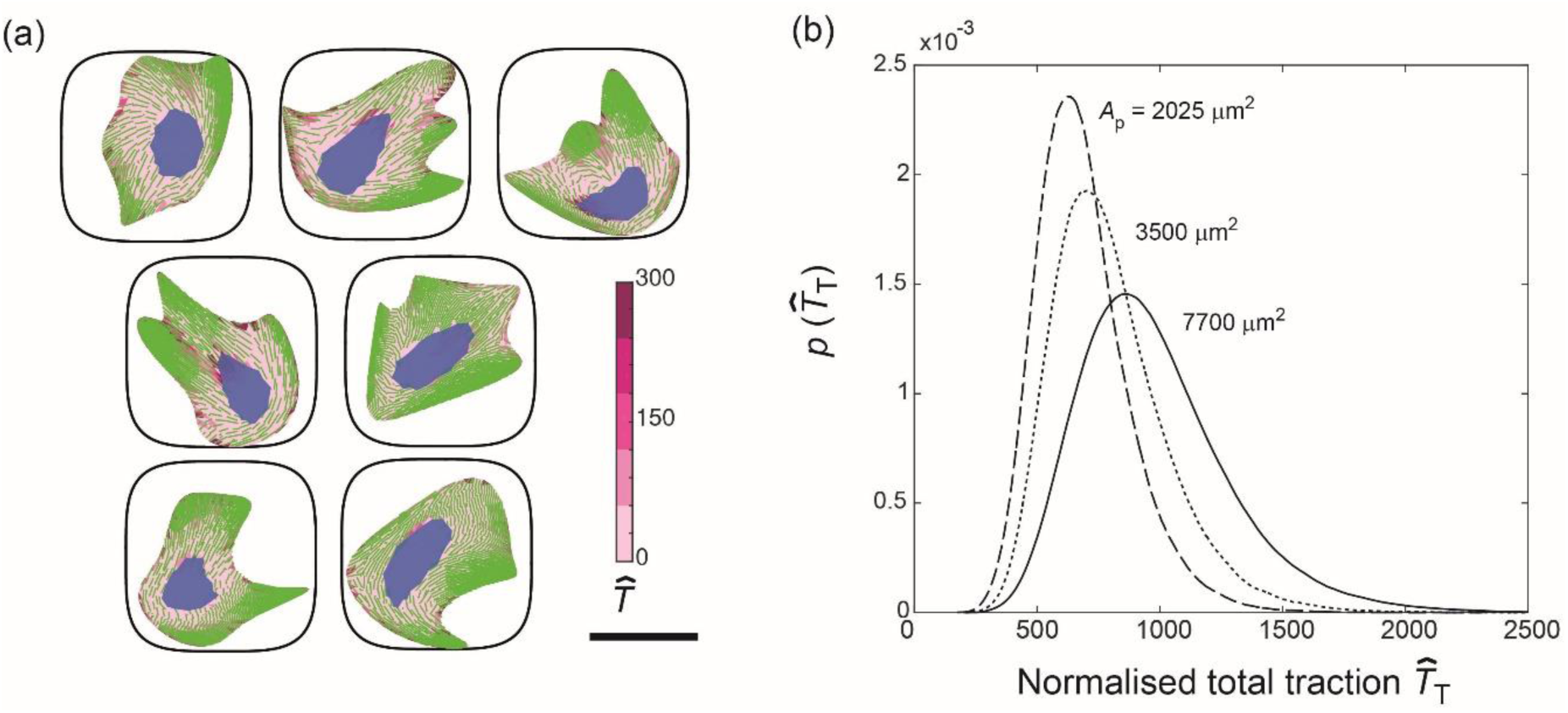
(a) A selection of 7 cell configurations selected from the homeostatic ensemble for hMSCs seeded on substrate patterned with adhesive islands of area *A*_P_ = 2025 μm^2^. The predictions are shown as combined immunofluorescence-like images showing stress-fibres (green), focal adhesions (pink) and nucleus (blue). Focal adhesions are parameterised by the magnitude of the normalised traction 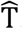. The scale bar = 30 μm. (b) Prediction of the probability density function of normalised total traction 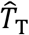 for hMSCs seeded on adhesive islands of selected areas *A*_P_.

### 3.3 Differentiation in growth media

In a landmark experiment, Engler et al. [6] established the influence of substrate elasticity on stem cell lineage commitment. Their experiments were performed in a growth medium which encouraged differentiation into osteoblasts, myoblasts and maybe adipocytes depending on substrate stiffness although the experiments reported in Engler et al [6] did not directly report commitment to adipocytes. Thus, here we only focus on the differentiation of hMSCs to osteoblasts, myoblasts and set 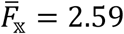 and 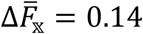 for osteoblasts while for myoblasts we set 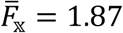 and 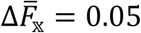 with *N*_c_ = 15.

Predictions of the fraction 𝒫_𝕩_ of the hMSCs differentiated into osteoblasts and myoblasts as a function of the substrate stiffness *E*_sub_ are included in Fig. 5a alongside measurements from Engler et al. [6]. Excellent agreement is obtained, with osteoblasts favoured on the stiffer substrates. We note that even for *E*_sub_ = 30 kPa where the probability of differentiation into osteoblasts peaks (and equally at *E*_sub_ = 10 kPa where the probability of differentiation into myoblasts is a maximum), the differentiation fraction 𝒫_𝕩_ ≠ 1, i.e. the model predicts that at *E*_sub_ = 30 kPa, approximately 20% of the hMSCs remain undifferentiated much like the measurements. The excellent agreement between predictions and observations is of course partially related to the fact that 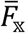 and 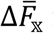 for myoblasts and osteoblasts are calibrated against the differentiation data of Engler et al. [6]. However, the strength of the approach is that the model can now be used to predict the differentiation response for hMSCs on adhesive islands and these predictions are included in Fig. 5b. While no data exists to-date for the differentiation of hMSCs in growth media seeded on adhesive islands, our model suggests a preference for osteoblasts on larger islands and myoblasts on smaller islands. The propensity for differentiation of hMSCs into osteoblasts on larger islands is known at-least for hMSCs cultured in mixed media (as will be discussed subsequently) but here our predictions suggest that the mechanical cues from substrate stiffness and geometric cues from sizes of adhesive islands can have similar effects on the differentiation response of hMSCs in growth media.

**Figure 5:**
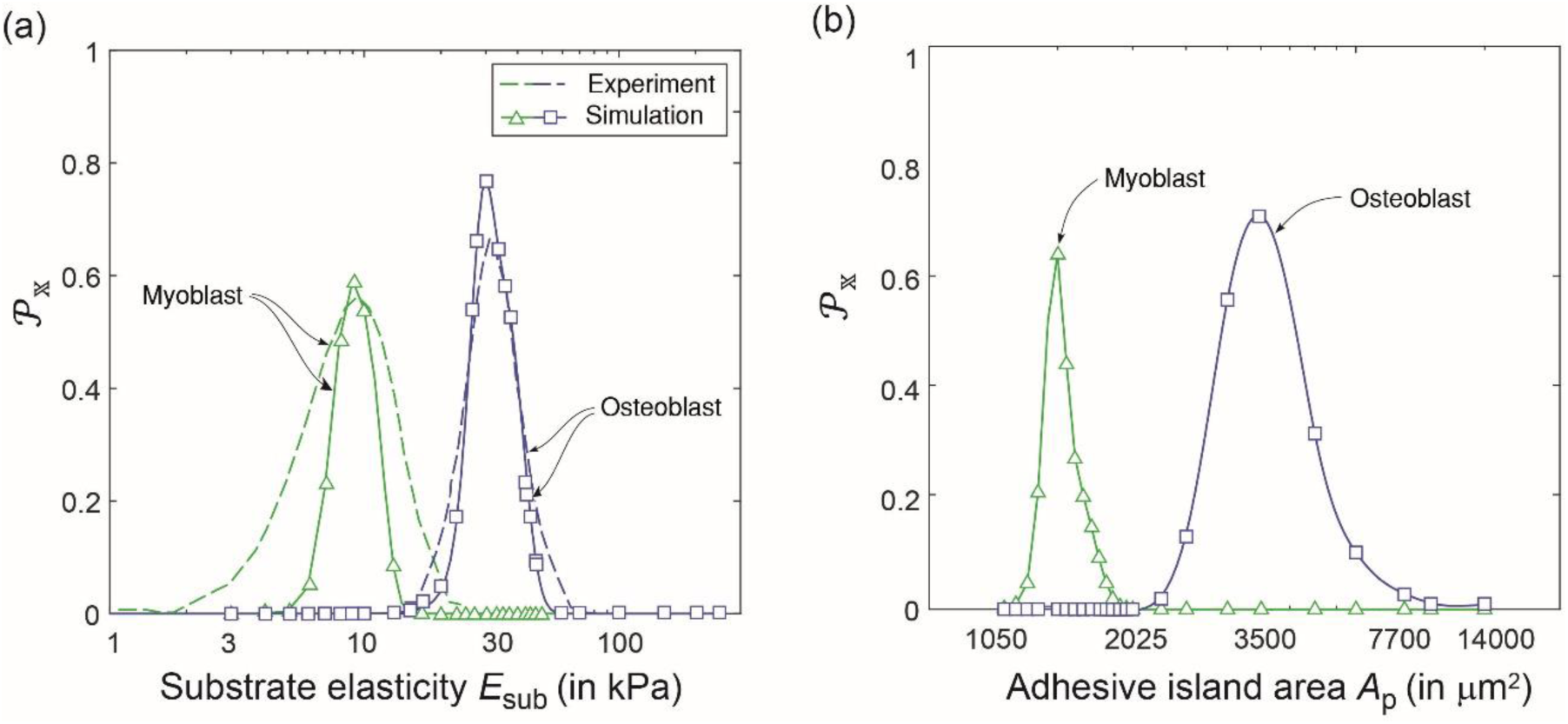
(a) Predictions of the variation of differentiation fraction 𝒫_𝕩_ for hMSCs seeded on elastic substrates with stiffness *E*_sub_ in growth media and compared with measurements from Engler et al. [6]. (b) Corresponding predictions of 𝒫_𝕩_ for hMSCs seeded on substrates patterns with adhesive islands of area *A*_P_ and seeded in growth media. The hMSCs in growth media are assumed to differentiate into osteoblasts, myoblasts or remain undifferentiated.

To understand the equivalency of these different cues, we first examine in further detail the differentiation predictions on elastic substrates. We have assumed that differentiation is set by the distribution of the cytoskeletal free-energy 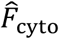. Predictions of the probability density functions of 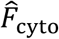 for hMSCs on substrates with three selected stiffness are included in Fig. 6a. As discussed in [26], higher substrate stiffness allows cells to exert larger tractions without a significant energy penalty, and the probability of cells to adopt configurations with larger cell areas, aspect ratios and higher levels of stress-fibre polymerisation increases. A direct consequence of the high level of stress-fibre polymerisation is lowering of the cytoskeletal free-energy as seen in Fig. 6a. These distributions then via (3) give the distribution of the cytoskeletal free-energy 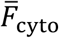 that cells assume over the 24-48 hour period after seeding during which they set their lineage. Predictions of the probability density functions of 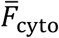 are included in Fig. 6b for the substrate stiffness employed in Fig. 6a. These distributions are relatively well dispersed suggesting that if the differentiation of hMSCs was set by this average cytoskeletal free-energy, their response on these three different substrates would vary substantially. In Fig. 6b, we have marked the range 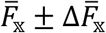 for the differentiation into osteoblasts in growth media. Clearly a large fraction of hMSCs seeded on the *E*_sub_ = 30 kPa will differentiate into osteoblasts as seen in Fig. 5a, with a very small fraction of cells on *E*_sub_ = 70 kPa also differentiating into osteoblasts as there is a small overlap in the distribution of 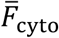 on *E*_sub_ = 70 kPa with the differentiation range 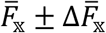 for osteoblasts. However, there is no overlap with hMSCs on *E*_sub_ = 10 kPa with no differentiation into osteoblasts expected for cells seeded on such soft substrates. Thus, the propensity of hMSCs to differentiate into osteoblasts when seeded on substrates with stiffness *E*_sub_ ≈ 30 kPa is directly related to the fact that their average cytoskeletal free-energy is in the correct range: for higher stiffness substrates 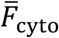 is too low due to the higher levels of stress-fibre polymerisation, while on softer substrates 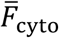 is too high due to the significantly reduced levels of stress-fibre polymerisation.

**Figure 6:**
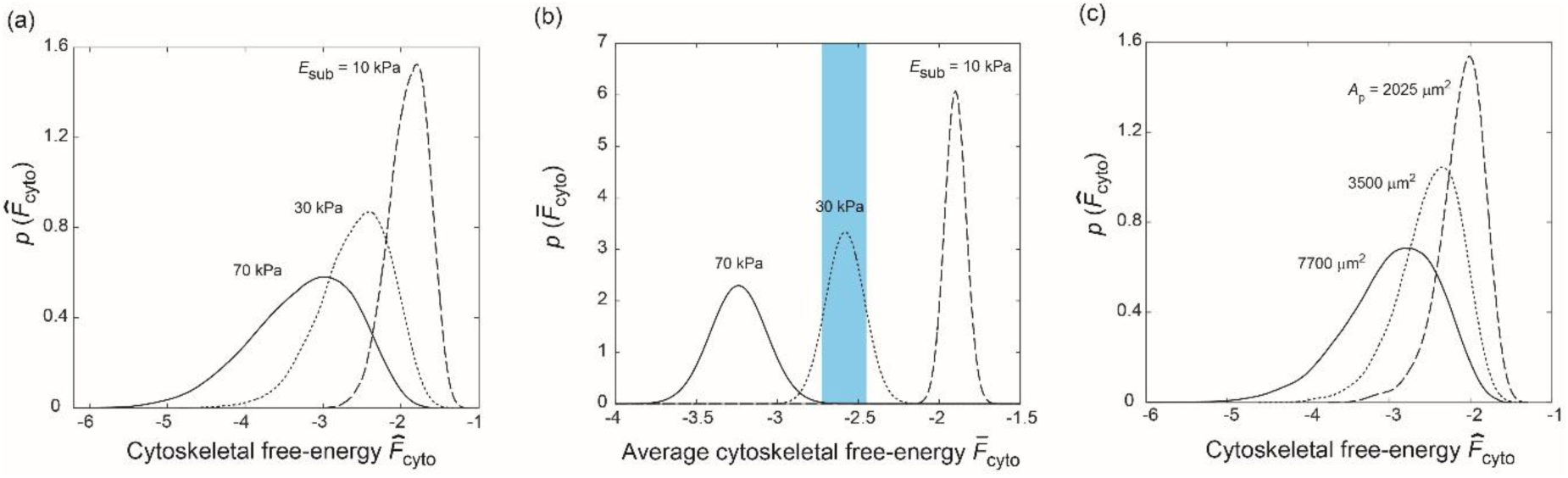
Predictions of the probability density function of (a) the cytoskeletal free-energy 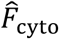 for hMSCs seeded on substrates of selected stiffness *E*_sub_, and (b) the corresponding distributions of the average cytoskeletal free-energy 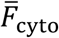. In (b), we have indicated the band (blue) of average cytoskeletal free-energies over which hMSCs are assumed to differentiate into osteoblasts in growth media. (c) Predictions of the probability density functions of 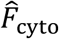 for hMSCs seeded on substrates patterned with adhesive islands of area *A*_P_.

Recall that with increasing substrate stiffness the cytoskeletal free-energy decreases, and this is associated with an increase in cell area. Seeding cells on rigid substrates patterned with adhesive islands can restrict the spreading of cells and thereby have a similar effect on the cytoskeletal free-energy by constraining stress-fibre polymerisation via this geometric cue rather than the stiffness cue. Predictions of the distribution of 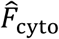 are included in Fig. 6c for selected adhesive island areas *A*_P_ with cytoskeletal free-energy again decreasing with increasing *A*_P_ (for *A*_P_ > 14000 μm^2^ the adhesive islands do not restrict cell spreading and the results converge to the *E*_sub_ = 70 kPa case discussed above). The consequences are therefore similar to the stiffness cues with hMSCs differentiating into osteoblasts for intermediate island areas.

### 3.4 Differentiation in mixed media

In mixed media, hMSCs differentiate into osteoblasts and adipocytes. We keep 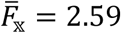 unchanged for osteoblasts and increase 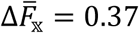 while we choose 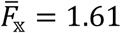 and 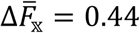 for adipocytes. These values are again chosen in order to obtain agreement with measurements for hMSCs seeded on adhesive islands in mixed media [3]. Predictions of the differentiation fraction 𝒫_𝕩_ for adipocytes and osteoblasts as a function of the area *A*_P_ of the adhesive islands (atop stiff substrates with stiffness *E*_sub_ = 70 kPa) and the corresponding predictions of 𝒫_𝕩_ as a function of substrate stiffness *E*_sub_ are included in Figs. 7a and 7b, respectively. The experimentally measured differentiation fraction from McBeath et al. [3] for an island size *A*_P_ = 2025 μm^2^ and from Guvendiren & Burdick [35] for cells cultured on substrates of stiffness *E*_sub_ = 3 kPa and 30 kPa included in Figs. 7a and 7b confirm the fidelity of the predictions. Importantly, the equivalency of the stiffness and geometric cues seen for growth media also carries forward to mixed media, where we now see an increased tendency for differentiation into adipocytes at either lower adhesive island areas or lower substrate stiffness.

**Figure 7:**
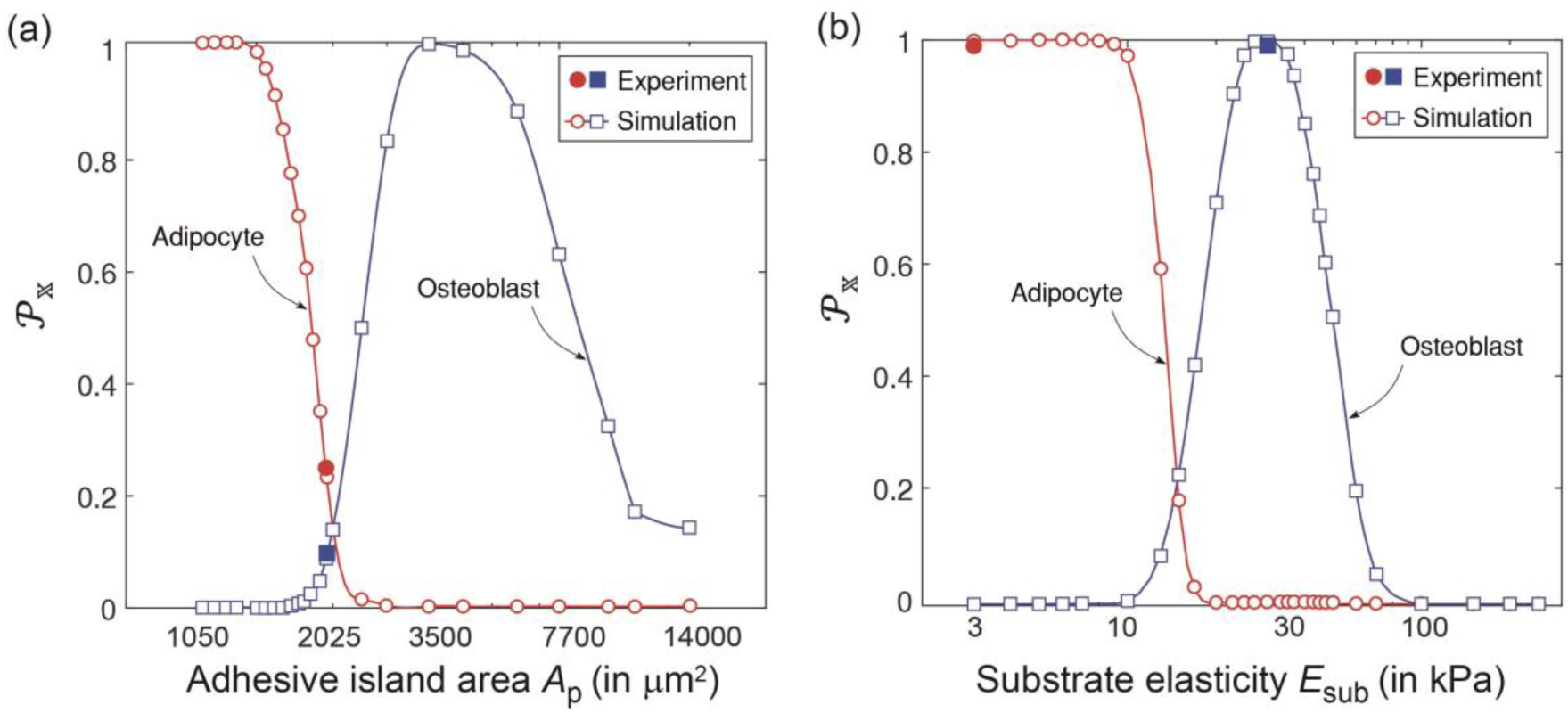
Predictions of the variation of differentiation fraction 𝒫_𝕩_ for hMSCs seeded on (a) substrates patterned with adhesive islands of area *A*_P_, and (b) elastic substrates with stiffness *E*_sub_ in mixed media. The hMSCs in mixed media are assumed to differentiate into osteoblasts, adipocytes or remain undifferentiated. Experimental measurements for an island of area *A*_P_ = 2025 μm^2^ from McBeath et al. [3] and substrates of stiffness *E*_sub_ = 3 kPa and 30 kPa from Guvendiren & Burdick [35] are included in (a) and (b), respectively.

Overall, the reason for this equivalency is as discussed above: adipocytes are favoured when the average cytoskeletal free-energy is higher and this occurs either by restricting cell spreading via the island size or on low stiffness substrates where the large tractions result in an energy penalty from the substrate which prevents cell spreading and enhances the cytoskeletal free-energy.

## Discussion

The equivalency of the cues discussed above, i.e. appropriately controlling adhesive island area can have an effect similar to substrate stiffness on hMSC differentiation seems to suggest that observable morphometrics such as cell area, aspect ratio etc. might correlate with the lineage the hMSCs adopt. In fact, it has been suggested [36] that changes to cell shape may be transduced into regulatory signals that govern cell fate. However, it is clearly seen from the distributions (Figs. 3a-c) that while cues such as substrate stiffness and island area do affect observables such as cell shape, the significant overlap of these distributions for the different cues strongly suggests that they cannot be directly used to determine cell fate. Here we claim that cytoskeletal free-energy, which gives a direct indication of the biochemical state of the cell, is a better metric to predict cell differentiation.

One way to directly show why simple observables are insufficient is to plot the predictions of the correlation between common observables (i.e., cell area *Â*, aspect ratio *A*_s_ and total traction 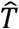) and the cytoskeletal free-energy 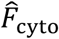. The joint-probability distributions of 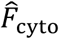 with each of these observables, i.e.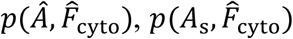 and 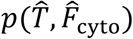 are shown in Figs. 8a-c for hMSCs seeded on an elastic substrate of stiffness *E*_sub_ = 30 kPa. These distributions are an alternative way of showing scatter plots typically used to judge correlation of variables with the yellow regions indicating regions of high probability (i.e. if observations were made, we would anticipate to obtain a large number of independent measurements in those regions) and dark blue indicating regions of low probability (i.e. we would expect that the probability of making a measurement in these regions is small). Clearly, there seems minimal correlation between these observables and 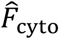 confirming our view that these direct observables are not adequate to predict hMSC commitment.

**Figure 8:**
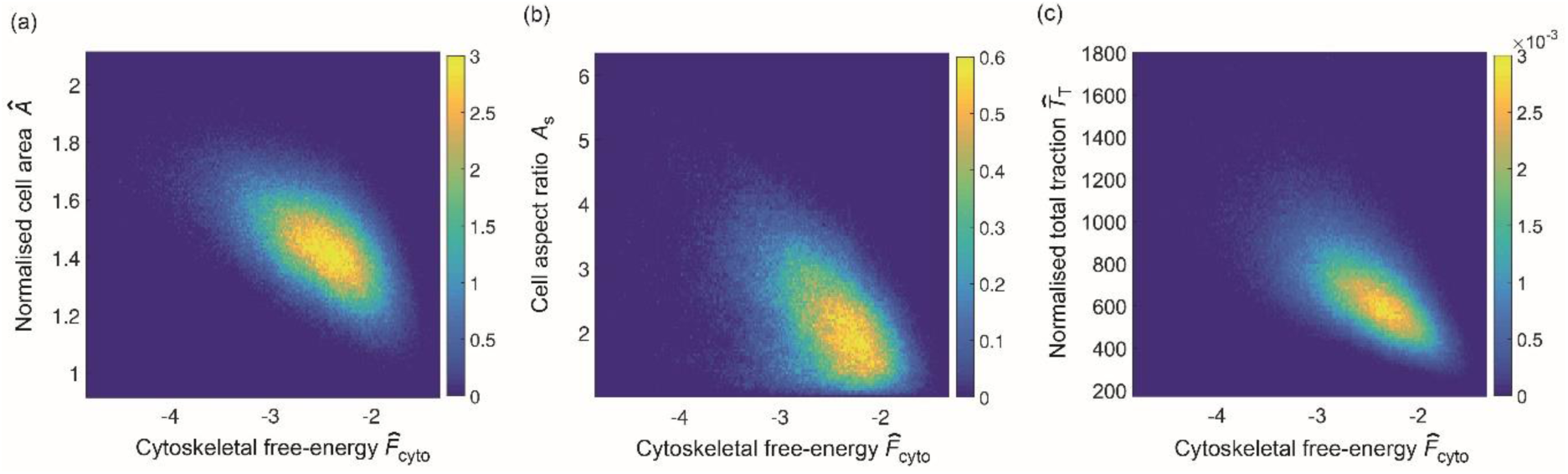
Predictions of the joint probability density distributions of the cytoskeletal free-energy 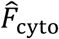 with the (a) normalised cell area *Â*, (b) cell aspect ratio *A*_s_ and (c) the normalised total traction 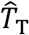. Results are shown for hMSCs seeded on an elastic substrate of stiffness *E*_sub_ = 30 kPa.

To further illustrate that direct observables such as cell shape or tractions may not be sufficient to predict cell differentiation, we computationally performed an inhibition study whereby we constrained cytoskeletal tension generation. This is an attempt to simulate drugs such as the Rho kinase (ROCK) inhibitor Y-27632 that restricts myosin activation. We simulated this by reducing the maximum tensile stress *σ*_max_ generated by a stress-fibre from 240 kPa to 231 kPa. Predictions of the distributions of *Â, A*_s_ and 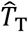 for cells cultured on a substrate patterned with adhesive islands of area *A*_P_ = 2725 μm^2^ are included in Figs. 9a-c for both the reference (untreated) case and with the simulated ROCK inhibitor. There is no appreciable difference in these direct observables. The corresponding predictions for the differentiation fraction 𝒫_𝕩_ for both the untreated and ROCK inhibitor treated cases are included in Fig. 10a. The untreated cells are predicted to show a strong commitment for osteoblasts, while the ROCK inhibitor treated cells are predicted to display a much weaker tendency to differentiate, and also predicted to be equally likely to differentiate into adipocytes and osteoblasts. These findings are consistent with the measurements of McBeath et al. [3], who performed drug inhibition studies specifically to experimentally test whether cell shape affects cell differentiation. In particular, they cultured hMSCs in mixed media in the presence of 10 μM Y-27632. Similar to our computational results, they observed that the treated cells remained spread and morphologically similar to the untreated cells but no longer exhibited the differentiation response of the untreated cells. Their measurements of the differentiation fractions for cells cultured on *A*_P_ = 2725 μm^2^ islands are included in Fig. 10a, and show excellent agreement with the computational results.

**Figure 9:**
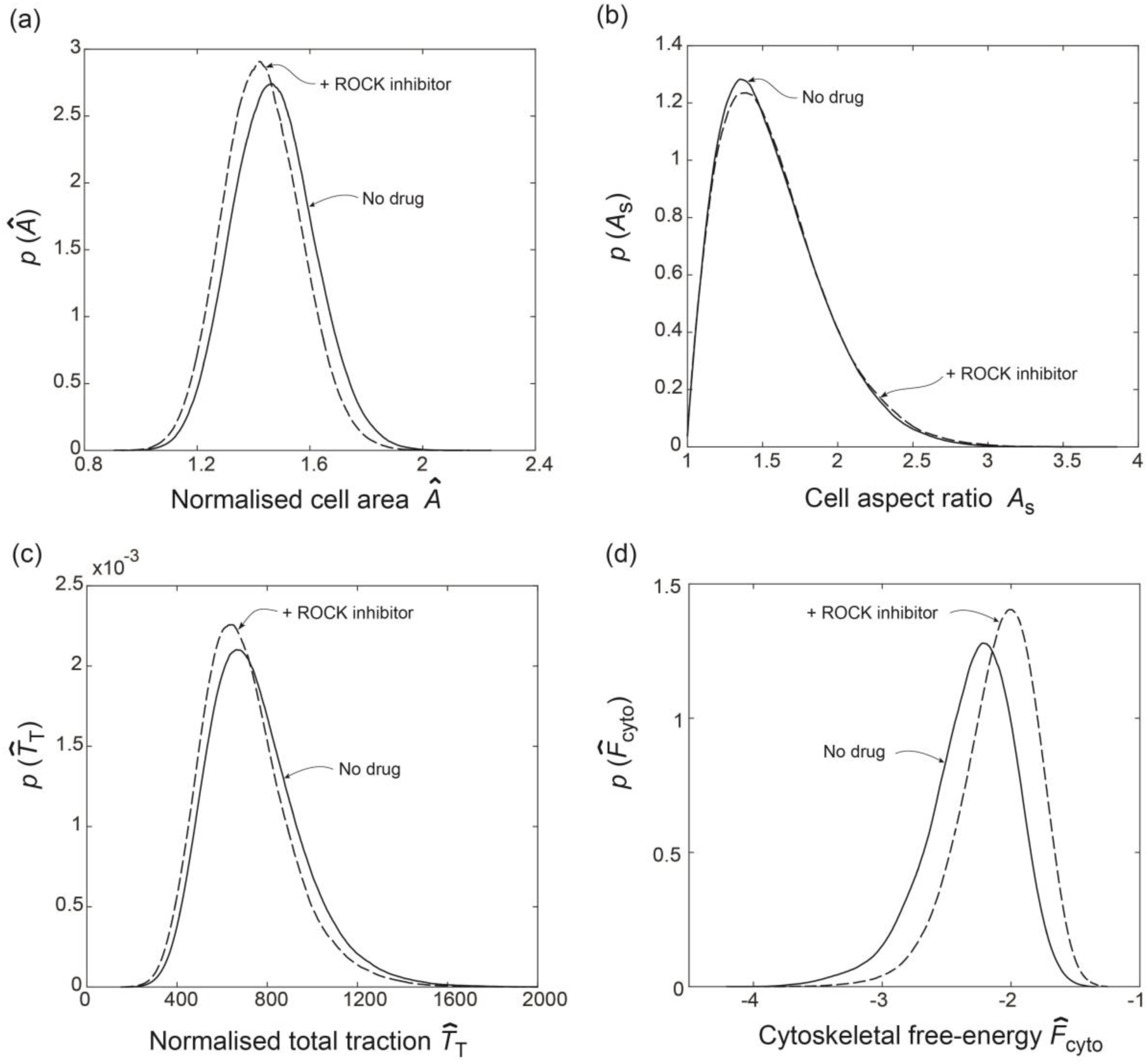
Predictions of the probability density functions for (a) normalised cell area *Â*, (b) cell aspect ratio *A*_s_, (c) normalised total traction 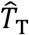, and (d) the cytoskeletal free-energy 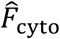 for hMSCs seeded on substrates patterned with adhesive islands of area *A*_p_ = 2725 μm^2^. Results are shown for both untreated cells and cells treated with a ROCK inhibitor.

**Figure 10:**
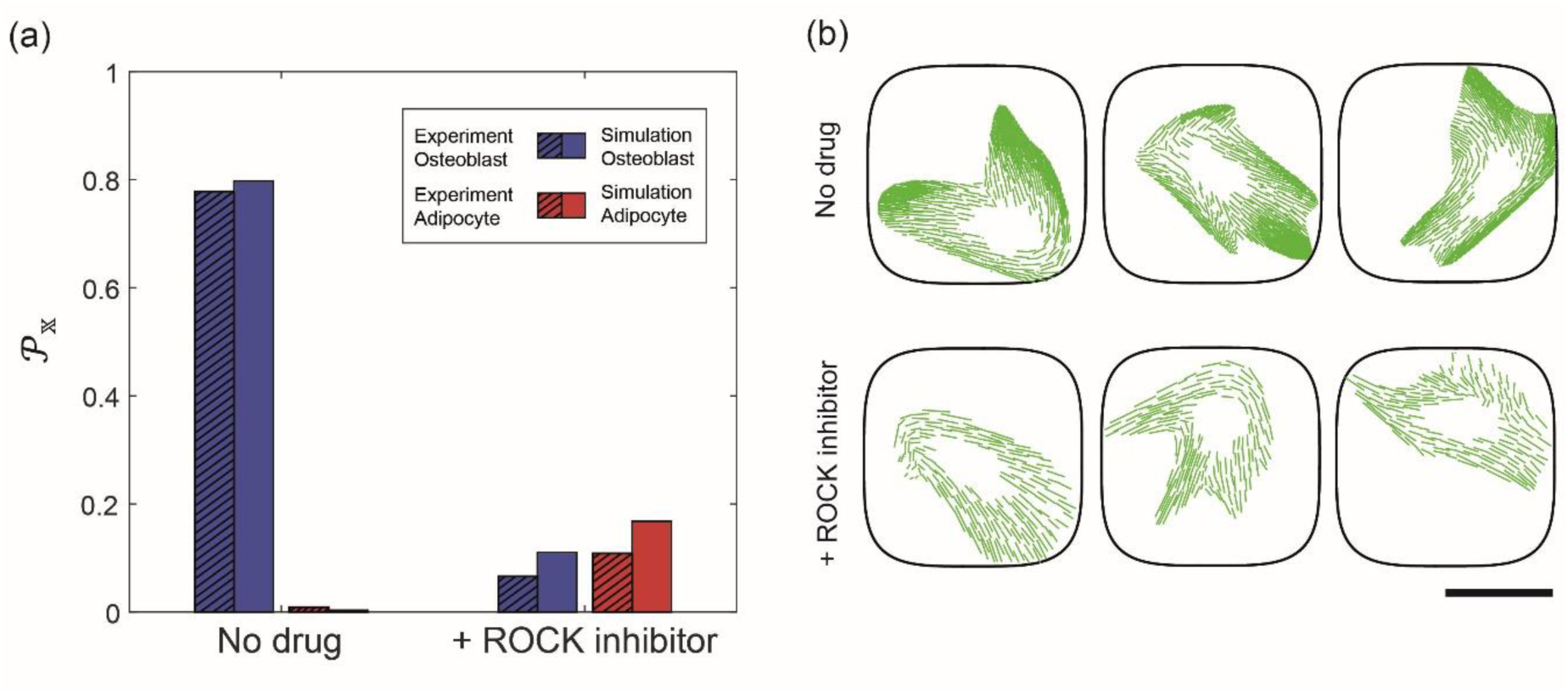
(a) Predictions of the differentiation fraction 𝒫_𝕩_ for hMSCs seeded on a substrate patterned with adhesive islands of area *A*_P_ = 2725 μm^2^. Results are shown for untreated cells as well as cells treated with a ROCK inhibitor, along with corresponding measurements from McBeath et al. [3] where Y-27632 was used as the ROCK inhibitor. (b) Three randomly selected cell morphologies from the entire homeostatic ensemble for untreated cells and cells treated with a ROCK inhibitor seeded on a substrate with *A*_P_ = 2725 μm^2^. The scale bar = 30 μm. In these images, we only show the stress-fibre distributions to illustrate the reduction in the level of stress-fibre polymerisation due to the treatment with a ROCK inhibitor.

The question arises as to why the computational model predicts such a dramatic change in the response given that common observables such as cell shape and tractions are seemingly unaffected when simulated with a ROCK inhibitor. Of course, differentiation in the model is directly related to the cytoskeletal free-energy 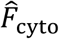. Simulating a ROCK inhibitor by reducing the value of *σ*_max_ from 240 kPa to 231 kPa does not affect the cell morphology and tractions substantially but does affect the state of the cytoskeleton and thereby 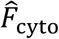, as seen in Fig 9d. In terms of direct observations, this will be seen as a reduction in the level of stress-fibre polymerisation in imaged cells. Three randomly selected cell morphologies (from the entire computed distribution of states the cells assume) for untreated and ROCK inhibitor treated cases are shown in Fig. 10b for hMSCs seeded on substrates with *A*_P_ = 2725 μm^2^. In these images only, the stress-fibre cytoskeleton has been marked in green similar to immunofluorescence imaging of actin in experiments. There is a clear reduction in the level of stress-fibre polymerisation due to the addition of the ROCK inhibitor in line with the observations of McBeath et al. [3]. It is this reduction in the level of stress-fibre polymerisation that enhances 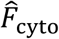 (see Fig. 9d) and inhibits cell differentiation even though there are no major changes in the cell morphometrics. The fact that no single cell morphometric is found to correlate with the lineage hMSCs adopt is well established with learning type models, in which using multiple metrics typically found to give reasonable predictions of cell fate [22, 23]. For example, it is conceivable that a combination of morphometrics such as cell area, aspect ratio and traction will correlate with cell fate for untreated cells. However, it is clear that for cell treated with a ROCK inhibitor the metrics will need to include a quantification of stress-fibre polymerisation. However, cytoskeletal free-energy which directly measures the biochemical state of the cell is a single metric that correlates with the lineages hMSCs adopt. Unfortunately, while 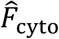 is not directly measurable in experiments, the fact that such a correlation might exist provides insight into the regulatory mechanisms that govern the commitment of hMSCs.

In summary, we hypothesise that the shape fluctuations of hMSCs in response to physical cues in their microenvironment during 1-2 days after seeding determine the probability and phenotype of cell lineage commitment and subsequent differentiation. The cell shape fluctuations are an output of the homeostatic mechanics framework, with the physical cues and a simple free-energy model for the cell as the only inputs. Analysed through the lens of a single biochemical parameter, i.e.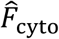, the aggregate of cell shape fluctuations in a given microenvironment can provide an early forecast of stem cell differentiation (within 1-2 days, compared to the typical forecast timescale of 1-2 weeks in experiments [3, 6]). The efficacy of the framework is demonstrated here under different conditions (mechanical and geometric cues, and in the presence of actomyosin inhibitors), thus inspiring confidence in its applicability to other insoluble cues in the stem cell niche. A key assumption in the free-energy model for the cell is that the morphological response of cells is independent of the culture medium. Changes to the free-energy model to account for the influence of culture medium in cellular morphological response can enhance the applicability of the homeostatic mechanics framework to predict differentiation response in the presence of soluble cues in the stem cell niche.

## Authors’ contributions

HS performed the simulations, processed the numerical data and created the figures. SSS, AV and HS contributed in developing the numerical codes for the model. VSD designed and supervised the project. VSD wrote the paper; all authors provided critical comments and edited the manuscript.

## References

1. Watt FM, Huck WTS. 2013 Role of the extracellular matrix in regulating stem cell fate. Nat. Rev. Mol. Cell Biol. 14, 467–473.

2. Sun Y, Chen CS, Fu J. 2012 Forcing Stem Cells to Behave: A Biophysical Perspective of the Cellular Microenvironment. Annu. Rev. Biophys. 41, 519–542.

3. McBeath R, Pirone DM, Nelson, CM, Bhadriraju K, Chen CS. 2004 Cell shape, cytoskeletal tension, and RhoA regulate stem cell lineage commitment. Dev. Cell 6, 483–495.

4. Discher DE, Mooney DJ, Zandstra PW. 2009 Growth factors, matrices, and forces combine and control stem cells. Science 324(5935): 1673–1677.

5. Zayzafoon M, Gathings WE, McDonald JM. 2004 Modeled microgravity inhibits osteogenic differentiation of human mesenchymal stem cells and increases adipogenesis. Endocrinology 145(5): 2421–2432.

6. Engler AJ, Sen S, Sweeney HL, Discher DE. 2006 Matrix elasticity directs stem cell lineage specification. Cell 126, 677–689.

7. Lee SJ, Yang S. 2017 Substrate curvature restricts spreading and induces differentiation of human mesenchymal stem cells. Biotechnol. J. 12, 1700360.

8. Fu J, Wang YK, Yang MT, Desai RA, Yu X, Liu Z, Chen CS. 2010 Mechanical regulation of cell function with geometrically modulated elastomeric substrates. Nature Methods 7, 733–736.

9. Sonam S, Sathe SR, Yim EKF, Sheetz MP, Lim CT. 2016 Cell contractility arising from topography and shear flow determines human mesenchymal stem cell fate. Sci. Rep. 6:20415.

10. Oh S, Brammer KS, Li YSJ, Teng D, Engler AJ, Chien S, Jin S. 2009 Stem cell fate dictated solely by altered nanotube dimension. Proc. Natl. Acad. Sci. USA, 106, 2130–2135.

11. Kim H, Kim I, Choi HJ, Kim SY, Yang EG. 2015 Neuron-like differentiation of mesenchymal stem cells on silicon nanowires. Nanoscale, 7, 17131–17138.

12. Teo BKK, Wong ST, Lim CK, Kung TYS, Yap CH, Ramagopal Y, Romer LH, Yim EKF. 2013 Nanotopography modulates mechanotransduction of stem cells and induces differentiation through focal adhesion kinase. ACS Nano, 7(6):4785–4798.

13. Ahn EH, Kim Y, Kshitiz, An SS, Afzal J, Lee S, Kwak M, Suh KY, Kim DH, Levchenko A. 2014 Spatial control of adult stem cell fate using nanotopographic cues. Biomaterials, 35, 2401–2410.

14. Dalby MJ, Gadegaard N, Tare R, Andar A, Riehle MO, Herzyk P, Wilkinson CDW, Oreffo ROC. 2007 The control of human mesenchymal cell differentiation using nanoscale symmetry and disorder. Nature Mater. 6, 997–1003.

15. Frith JE, Mills RJ, Cooper-White JJ. 2011 Lateral spacing of adhesion peptides influences human mesenchymal stem cell behaviour. J. Cell Sci. 125, 317–327.

16. Kilian KA, Bugarija B, Lahn BT, Mrksich M. 2010 Geometric cues for directing the differentiation of mesenchymal stem cells. Proc. Natl Acad. Sci. USA 107, 4872–4877.

17. Heydari T, Heidari M, Mashinchian O, Wojcik M, Xu K, Dalby MJ, Mahmoudi M, Ejtehadi MR. 2017 Development of a virtual cell model to predict cell response to substrate topography. ACS Nano, 11, 9084–9092.

18. Mousavi SJ, Doweidar MH. 2015 Role of mechanical cues in cell differentiation and proliferation: a 3D numerical model. PLoS ONE, 10:e0124529.

19. Macis M, Lugli F, Zerbetto F. 2017 Modeling living cells response to surface tension and chemical patterns. ACS Appl. Mater. Interfaces, 9, 19552–19561.

20. Deshpande RS, Spector AA. 2017 Modeling stem cell myogenic differentiation. Sci. Rep. 7:40639.

21. Peng T, Liu L, MacLean AL, Wong CW, Zhao W, Nie Q. 2017 A mathematical model of mechanotransduction reveals how mechanical memory regulates mesenchymal stem cell fate decisions. BMC Syst. Biol., 11:55.

22. Treiser MD, Yang EH, Gordonov S, Cohen DM, Androulakis IP, Kohn J, Chen CS, Moghe PV. 2009 Cytoskeleton-based forecasting of stem cell lineage fates. Proc. Natl. Acad. Sci. USA, 107, 610–615.

23. Cutiongco MFA, Jensen BS, Reynolds PM, Gadegaard N. 2019 Predicting gene expression using morphological cell responses to nanotopography. bioRxiv, https://doi.org/10.1101/495879

24. Stumpf PS, Smith RCG, Lenz M, Schuppert A, Müller F-J, Babtie A, et al. 2017 Stem cell differentiation as a non-Markov stochastic process. Cell Syst., 5, 268–282.

25. Kalmar T, Lim C, Hayward P, Muñoz-Descalzo S, Nichols J, Garcia-Ojalvo J, Arias AM. 2009 Regulated fluctuations in nanog expression mediate cell fate decisions in embryonic stem cells. PLoS Biol., 7:e1000149.

26. Shishvan SS, Vigliotti A, Deshpande VS. 2018 The homeostatic ensemble for cells. Biomech Model Mechanobiol, 17(6):1631–1662.

27. McEvoy E, Shishvan SS, Deshpande VS, McGarry JP. 2018 Thermodynamic modeling of the statistics of cell spreading on ligand-coated elastic substrates. Biophys. J. 115, 2451–2460.

28. Buskermolen ABC, Suresh H, Shishvan SS, Vigliotti A, DeSimone A, Kurniawan NA, Bouten CVC, Deshpande VS. 2019 Entropic forces drive cellular contact guidance. Biophys. J., 116, 1994–2008.

29. Recordati G, Bellini TG. 2004 A definition of internal constancy and homeostasis in the context of non-equilibrium thermodynamics. Exp. Physiol. 89, 27–38.

30. Weiss TF. 1996 Cellular biophysics. MIT Press, Cambridge.

31. Vigliotti A, Ronan W, Baaijens FPT, Deshpande VS. 2015 A thermodynamically motivated model for stress-fibre reorganization. Biomech. model. Mechanobiol. 15:761–789.

32. Ronan W, Deshpande VS, McMeeking RM, McGarry JP. 2014 Cellular contractility and substrate elasticity: a numerical investigation of the actin cytoskeleton and cell adhesion. Biomech. Model. Mechanobiol, 13, 417–435.

33. Vernerey FJ, Farsad M. 2014 A mathematical model of the coupled mechanisms of cell adhesion, contraction and spreading. J. Math. Biol, 68, 989–1022.

34. Kimmel JC, Chang AY, Brack AS, Marshall WF. 2018 Inferring cell state by quantitative motility analysis reveals a dynamic state system and broken detailed balance. PLoS Comput. Biol., 14:e1005927.

35. Guvendiren M, Burdick JA. 2012 Stiffening hydrogels to probe short- and long-term cellular responses to dynamic mechanics. Nat. Commun. 3:792.

36. Huang S, Chen CS, Ingber DE. 1998 Control of cyclin D1, p27(Kip1), and cell cycle progression in human capillary endothelial cells by cell shape and cytoskeletal tension. Mol. Biol. Cell, 9, 3179–3193.

