## Supplementary Information for "Free-energy-based framework for early forecasting of stem cell differentiation"

**Supplementary Information 1: Gibbs free-energy of a morphological microstate**

Here, we provide a brief overview of the Gibbs free-energy model for morphological microstates required by the homeostatic mechanics framework of Shishvan et al. [1]. The aim is to provide the reader the key aspects of the free-energy model required for fully appreciating the computational results presented in the main text. Readers are referred to Shishvan et al. [1] for a more complete treatment including the derivations of the relevant equations.

### *1.1 The equilibrium Gibbs free-energy of a morphological microstate*

Similar to conventional statistical mechanics calculations that require a model for the energy of the system, the homeostatic statistical mechanics framework requires a model for the Gibbs free-energy $G^{(j)}$ of morphological microstate $(j)$. Here, we calculate $G^{(j)}$ using the free-energy model of Vigliotti et al. [2], as modified in [1] and [3], that includes contributions from cell elasticity and the actin/myosin stress-fibre cytoskeleton, with the cell modelled as a two-dimensional (2D) body in the $x_1 - x_2$ plane adhered to an elastic substrate (Fig. 1b of the main text) such that the out-of-plane Cauchy stress $\Sigma_{33} = 0$. The state of the system changes as the cell moves, spreads and changes shape on the elastic substrate. Here, we shall give a prescription to calculate the Gibbs free-energy of the system when the cell is in a specific morphological microstate $(j)$, where the connections of material points on the cell membrane to the surface of the substrate are specified. Calculating the Gibbs free-energy of the system when the material points on the cell membrane is connected to the surface of the adhesive island requires a small modification to the prescription outlined below. Readers are urged to refer to Supplementary S1.2 (and also Supplementary S2 of Buskermolen et al [3]) for further information. In the following, for the sake of notational brevity, we shall drop the superscript $(j)$ that denotes the morphological microstate, as the entire discussion refers to a single morphological microstate.

The Vigliotti et al. [2] model assumes only two components within the cell: (i) a passive elastic contribution from subcellular structures such as the cell membrane, intermediate filaments and microtubules, and (ii) an active contribution from contractile acto-myosin stress-fibres that are modelled explicitly with the nucleus not explicitly modelled. This model was modified in [1] to incorporate a non-dilute concentration of stress-fibres, and further modified in [3] to include the nucleus in the analysis as a passive elastic body, in addition to the cytoplasm comprising the two above mentioned components. We shall first describe the modelling of the active acto-myosin stress-fibres in the cytoplasm and then discuss the elastic model of both the nucleus and the cytoplasm.

Consider a two-dimensional (2D) cell of thickness $b_0$ and volume $V_0$ in its elastic resting state comprising a nucleus of volume $V_{\mathrm{N}}$ and cytoplasm of volume $V_{\mathrm{C}}$ such that $V_0 = V_{\mathrm{N}} + V_{\mathrm{C}}$ (Fig. 1b of the main text). The representative volume element (RVE) of the stress-fibres within the cytoplasm in this resting configuration is assumed to be a cylinder of volume $V_{\mathrm{R}} = \pi b_0 \left(\frac{n^{\mathrm{R}}\ell_0}{2}\right)^2$, where $\ell_0$ is the length of a stress-fibre functional unit in its ground-state, and $n^{\mathrm{R}}$

is the number of these ground-state functional units within this reference RVE. The total number of functional unit packets within the cell is $N_0^{\mathrm{T}}$, and we introduce $N_0 = N_0^{\mathrm{T}} V_{\mathrm{R}}/V_{\mathrm{C}}$ as the average number of functional unit packets available per RVE; $N_0$ shall serve as a useful normalisation parameter. The state of stress-fibres at location $x_i$ within the cell is described by their angular concentration $\eta(x_i, \varphi)$, and there are $n(x_i, \varphi)$ functional units in series along the length of each stress-fibre in the RVE. Here, $\varphi$ is the angle of the stress-fibre bundle in the undeformed configuration with respect to the $x_2 -$ direction of the elastic substrate (Fig. 1b of the main text). Vigliotti et al. [2] showed that, at steady-state, the number $n^{\mathrm{ss}}$ of functional units within the stress-fibres is given by

$$\hat{n}^{\mathrm{ss}} \equiv \frac{n^{\mathrm{ss}}}{n^{\mathrm{R}}} = \frac{[1 + \varepsilon_{\mathrm{nom}}(x_i, \varphi)]}{1 + \tilde{\varepsilon}_{\mathrm{nom}}^{\mathrm{ss}}}, \tag{1.1}$$

where $\tilde{\varepsilon}_{\mathrm{nom}}^{\mathrm{ss}}$ is the strain at steady-state within a functional unit of the stress-fibres, and $\varepsilon_{\mathrm{nom}}(x_i, \varphi)$ is the nominal strain in direction $\varphi$. The chemical potential of the functional units within the stress-fibres in terms of the Boltzmann constant $k_{\mathrm{B}}$ is given by [1]

$$\chi_{\mathrm{b}} = \frac{\mu_{\mathrm{b}}}{n^{\mathrm{R}}} + k_{\mathrm{B}} T \ln\left[\left(\frac{\pi\, \hat{\eta}\, \hat{n}^{\mathrm{ss}}}{\hat{N}_{\mathrm{u}}\left(1 - \frac{\hat{\eta}}{\hat{\eta}_{\mathrm{max}}}\right)}\right)^{\frac{1}{n^{\mathrm{ss}}}} \left(\frac{\hat{N}_{\mathrm{u}}}{\pi \hat{N}_{\mathrm{L}}}\right)\right], \tag{1.2}$$

where the normalized concentration of the unbound stress-fibre proteins is $\hat{N}_{\mathrm{u}} \equiv N_{\mathrm{u}}/N_0$ with $\hat{\eta} \equiv \eta n^{\mathrm{R}}/N_0$, while $\hat{\eta}_{\mathrm{max}}$ is the maximum normalised value of $\hat{\eta}$ corresponding to full occupancy of all available sites for stress-fibres (in a specific direction) and $\hat{N}_{\mathrm{L}}$ is the number of lattice sites available to unbound proteins. The enthalpy $\mu_{\mathrm{b}}$ of $n^{\mathrm{R}}$ bound functional units at steady-state is given in terms of the isometric stress-fibre stress $\sigma_{\mathrm{max}}$ and the internal energy $\mu_{\mathrm{b0}}$ as

$$\mu_{\mathrm{b}} = \mu_{\mathrm{b0}} - \sigma_{\mathrm{max}} \Omega (1 + \tilde{\varepsilon}_{\mathrm{nom}}^{\mathrm{ss}}), \tag{1.3}$$

where $\Omega$ is the volume of $n^{\mathrm{R}}$ functional units. By contrast, the chemical potential of the unbound proteins is independent of stress and given in terms of the internal energy $\mu_{\mathrm{u}}$ as

$$\chi_{\mathrm{u}} = \frac{\mu_{\mathrm{u}}}{n^{\mathrm{R}}} + k_{\mathrm{B}} T \ln\left(\frac{\hat{N}_{\mathrm{u}}}{\pi\, \hat{N}_{\mathrm{L}}}\right). \tag{1.4}$$

For a fixed configuration of the 2D cell (i.e. a fixed strain distribution $\varepsilon_{\mathrm{nom}}(x_i, \varphi)$), the contribution to the specific Helmholtz free-energy of the cell $f$ from the stress-fibre cytoskeleton follows as

$$f_{\mathrm{cyto}} = \rho_0 \left(\hat{N}_{\mathrm{u}} \chi_{\mathrm{u}} + \int_{-\pi/2}^{\pi/2} \hat{\eta}\, \hat{n}^{\mathrm{ss}} \chi_{\mathrm{b}} d\varphi\right), \tag{1.5}$$

where $\rho_0 \equiv N_0/V_{\mathrm{R}}$ is the number of protein packets per unit reference volume available to form functional units in the cell. However, we cannot yet evaluate $f_{\mathrm{cyto}}$ as $\hat{N}_{\mathrm{u}}(x_i)$ and $\hat{\eta}(x_i, \varphi)$ are unknown. These will follow from the chemical equilibrium of the cell as will be discussed in Section 1.1.1.

The total stress $\Sigma_{ij}$ within the cell includes contributions from the passive elasticity provided mainly by the intermediate filaments of the cytoskeleton attached to the nuclear and plasma membranes and the microtubules, as well as the active contractile stresses of the stress-fibres. The total Cauchy stress is written in an additive decomposition as

$$\Sigma_{ij} = \sigma_{ij} + \sigma_{ij}^{\mathrm{p}}, \tag{1.6}$$

where $\sigma_{ij}$ and $\sigma_{ij}^{\mathrm{p}}$ are the active and passive Cauchy stresses, respectively. In the 2D setting with the cell lying in the $x_1 - x_2$ plane, the active stress is given in terms of the volume fraction $\mathcal{H}_0$ of the stress-fibre proteins as

$$\begin{bmatrix} \sigma_{11} & \sigma_{12} \\ \sigma_{12} & \sigma_{22} \end{bmatrix} = \frac{\mathcal{H}_0 \sigma_{\max}}{2} \int_{-\pi/2}^{\pi/2} \hat{\eta}[1 + \varepsilon_{\mathrm{nom}}(\varphi)] \begin{bmatrix} 2\sin^2\varphi^* & -\sin 2\varphi^* \\ -\sin 2\varphi^* & 2\cos^2\varphi^* \end{bmatrix} d\varphi, \tag{1.7}$$

where $\varphi^*$ is the angle of the stress-fibre measured with respect to $x_2$, and is related to its orientation $\varphi$ in the undeformed configuration by the rotation with respect to the undeformed configuration. The passive elasticity in the 2D setting is given by a 2D specialization of the Ogden [4] hyperelastic model as derived in [1]. The strain energy density function of this 2D Ogden model is

$$\Phi_{\mathrm{C}} \equiv \frac{2\mu_{\mathrm{C}}}{m_{\mathrm{C}}^2} \left[ \left(\frac{\lambda_{\mathrm{I}}}{\lambda_{\mathrm{II}}}\right)^{\frac{m_{\mathrm{C}}}{2}} + \left(\frac{\lambda_{\mathrm{II}}}{\lambda_{\mathrm{I}}}\right)^{\frac{m_{\mathrm{C}}}{2}} - 2 \right] + \frac{\kappa_{\mathrm{C}}}{2} (\lambda_{\mathrm{I}}\lambda_{\mathrm{II}} - 1)^2, \tag{1.8}$$

for the cytoplasm and

$$\Phi_{\mathrm{N}} \equiv \frac{2\mu_{\mathrm{N}}}{m_{\mathrm{N}}^2} \left[ \left(\frac{\lambda_{\mathrm{I}}}{\lambda_{\mathrm{II}}}\right)^{\frac{m_{\mathrm{N}}}{2}} + \left(\frac{\lambda_{\mathrm{II}}}{\lambda_{\mathrm{I}}}\right)^{\frac{m_{\mathrm{N}}}{2}} - 2 \right] + \frac{\kappa_{\mathrm{N}}}{2} (\lambda_{\mathrm{I}}\lambda_{\mathrm{II}} - 1)^2, \tag{1.9}$$

for the nucleus where $\lambda_{\mathrm{I}}$ and $\lambda_{\mathrm{II}}$ are the principal stretches, $\mu_{\mathrm{C}}$ ($\mu_{\mathrm{N}}$) and $\kappa_{\mathrm{C}}$ ($\kappa_{\mathrm{N}}$) the shear modulus and in-plane bulk modulus of cytoplasm (nucleus), respectively, while $m_{\mathrm{C}}$ ($m_{\mathrm{N}}$) is a material constant governing the non-linearity of the deviatoric elastic response of cytoplasm (nucleus). The cell is assumed to be incompressible, and thus throughout the cell, we set the principal stretch in the $x_3$ −direction $\lambda_{\mathrm{III}} = 1/(\lambda_{\mathrm{I}}\lambda_{\mathrm{II}})$. The (passive) Cauchy stress then follows as $\sigma_{ij}^{\mathrm{p}} p_j^{(k)} = \sigma_k^{\mathrm{p}} p_i^{(k)}$ in terms of the principal (passive) Cauchy stresses $\sigma_k^{\mathrm{p}}$ ($\equiv \lambda_k \partial\Phi_{\mathrm{C}}/\partial\lambda_k$ for the cytoplasm and $\equiv \lambda_k \partial\Phi_{\mathrm{N}}/\partial\lambda_k$ for the nucleus) and the unit vectors $p_j^{(k)}$ ($k = \mathrm{I}, \mathrm{II}$) denoting the principal directions. The total specific Helmholtz free-energy of the cell is then $f = f_{\mathrm{cyto}} + \Phi_{\mathrm{C}}$ in the cytoplasm and $f = \Phi_{\mathrm{N}}$ in the nucleus.

#### *1.1.1 Equilibrium of the morphological microstate*

Shishvan et al. [1] have shown that equilibrium of a morphological microstate reduces to two conditions: (i) mechanical equilibrium with $\Sigma_{ij,j} = 0$ throughout the system, and (ii) chemical equilibrium such that $\chi_{\mathrm{u}}(x_i) = \chi_{\mathrm{b}}(x_i, \varphi) = \text{constant}$, i.e. the chemical potentials of bound and unbound stress-fibre proteins are equal throughout the cell.

The condition $\chi_\mathrm{u} = \chi_\mathrm{b}$ implies that $\hat{\eta}(x_i, \varphi)$ is given in terms of $\hat{N}_\mathrm{u}$ by

$$\hat{\eta}(x_i, \varphi) = \frac{\hat{N}_\mathrm{u}\, \hat{\eta}_\mathrm{max} \exp\left[\frac{\hat{n}^\mathrm{ss}(\mu_\mathrm{u} - \mu_\mathrm{b})}{k_\mathrm{B} T}\right]}{\pi \hat{n}^\mathrm{ss} \hat{\eta}_\mathrm{max} + \hat{N}_\mathrm{u} \exp\left[\frac{\hat{n}^\mathrm{ss}(\mu_\mathrm{u} - \mu_\mathrm{b})}{k_\mathrm{B} T}\right]}, \tag{1.10}$$

and $\hat{N}_\mathrm{u}$ follows from the conservation of stress-fibre proteins throughout the cytoplasm, viz.

$$\hat{N}_\mathrm{u} + \frac{1}{V_\mathrm{C}} \int_{V_\mathrm{C}} \int_{-\pi/2}^{\pi/2} \hat{\eta}\, \hat{n}^\mathrm{ss} d\varphi \; dV = 1. \tag{1.11}$$

Knowing $\hat{N}_\mathrm{u}$ and $\hat{\eta}(x_i, \varphi)$, the stress $\Sigma_{ij}$ can now be evaluated and these stresses within the system (i.e. cell and substrate) need to satisfy mechanical equilibrium, i.e. $\Sigma_{ij,j} = 0$. In this case, the mechanical equilibrium condition is readily satisfied as the stress field $\Sigma_{ij}$ within the cell is equilibrated by a traction field $\mathrm{T}_i$ exerted by the substrate on the cell such that $b\Sigma_{ij,j} = -\mathrm{T}_i$, where $b(x_i)$ is the thickness of the cell in the current configuration.

The equilibrium value of the Gibbs free-energy is then given as $G = F_\mathrm{cell} + F_\mathrm{sub}$ where

$$F_\mathrm{cell} \equiv \rho_0 V_\mathrm{C} \chi_\mathrm{u} + \int_{V_\mathrm{C}} \Phi_\mathrm{C}\, dV + \int_{V_\mathrm{N}} \Phi_\mathrm{N}\, dV, \tag{1.12}$$

and

$$F_\mathrm{sub} \equiv \int_{V_\mathrm{sub}} \psi\, dV, \tag{1.13}$$

where ψ is the strain-energy density of the substrate material, and $V_\mathrm{sub}$ is the volume of the substrate. Here, $\chi_\mathrm{u}$ is given by Eq. (1.4) with the equilibrium value of $\hat{N}_\mathrm{u}$ obtained from Eq. (1.11). Details of numerical computation of $F_\mathrm{sub}$ is provided in [1]. For the purposes of further discussion, we label the equilibrium value $F_\mathrm{cyto} \equiv \rho_0 V_\mathrm{C} \chi_\mathrm{u}$ as the cytoskeletal free-energy of the cell, $F_\mathrm{passive} \equiv \int_{V_\mathrm{C}} \Phi_\mathrm{C}\, dV + \int_{V_\mathrm{N}} \Phi_\mathrm{N}\, dV$ as the passive elastic energy of the cell, and $F_\mathrm{sub}$ as the substrate free-energy.

#### *1.2 Numerical methods*

We employ Markov Chain Monte Carlo (MCMC) to construct a Markov chain that is representative of the homeostatic ensemble. This involves three steps: (i) a discretization scheme to represent morphological microstate ($j$), (ii) calculation of $G^{(j)}$ for a given morphological microstate ($j$), and (iii) construction of a Markov chain comprising these morphological microstates. Here, we briefly describe the procedure which was implemented in MATLAB.

In the general setting of a three-dimensional (3D) cell, a morphological microstate is defined by the connection of material points on the cell membrane to the surface of the substrate. In the 2D context of cells on micropatterned substrates, this reduces to specifying the connection of all material points of the cell to locations within the elastic substrate, i.e. a displacement field $u_i^{(j)}(X_i)$ is imposed on the cell with $X_i$ denoting the location of material points on the cell in the undeformed configuration, and these are then displaced to $x_i^{(j)} = X_i + u_i^{(j)}$ in morphological microstate $(j)$, such that all $x_i^{(j)}$ lie within the elastic substrate. These material points located at $x_i^{(j)}$ are then connected to material points on the substrate at the same location $x_i^{(j)}$, completing the definition of the morphological microstate in the 2D setting.

The cell is modelled as a continuum and thus $u_i^{(j)}$ is a continuous field. To calculate the density of the morphological microstates, we define $u_i^{(j)}$ via Non-Uniform Rational B-splines (NURBS) such that the morphological microstate is now defined by $M$ pairs of weights $\left[U_L^{(j)}, V_L^{(j)}\right]$ $(L = 1, \ldots, M)$. In all the numerical results presented here, we employ $M = 16$ with $4 \times 4$ weights $U_L^{(j)}$ and $V_L^{(j)}$ governing the displacements in the $x_1$ and $x_2$ directions, respectively. The NURBS employ fourth order base functions for both the $x_1$ and $x_2$ directions, and the knots vector included two nodes each with multiplicity four, located at the extrema of the interval. We emphasise here that this choice of representing the morphological microstates imposes restrictions on the morphological microstates that will be considered. Therefore, the choice of the discretization used to represent $u_i^{(j)}$ needs to be chosen so as to be able to represent the microstates we wish to sample, e.g. the choice can be based on the minimum width of a filopodium one expects for the given cell type. Given $u_i^{(j)}$, we can calculate $G^{(j)}$ using the model outlined in the main paper with the cell discretised using constant strain triangles of size $e \approx R_0/10$, where $R_0$ is the radius of the cell in its undeformed configuration.

We construct, via MCMC, a Markov chain that serves as a sample of the homeostatic ensemble for cells on wide elastic substrates and cells confined to square adhesive islands of area $A_{\mathrm{p}}$. The algorithm closely follows the approach developed by Shishvan et al. [1]. However, here we additionally model cells constrained on adhesive islands with cell adhesion outside the islands prevented. Over the range of island sizes used in the experimental investigation, cells that were partially or entirely outside the islands were not observed. Thus, we construct a sample of the homeostatic ensemble comprising solely of morphological microstates that are fully within the adhesive islands. This is done using the Metropolis [5] algorithm in an iterative manner using the procedure explained in detail in [1] (see *section 4.3* therein) but now with the following modification. In constructing the Markov chain, if any portion of the proposed new configuration of the cell lies outside the adhesive island, then the nodal boundary points outside the island were pushed back to the boundary of the island along a line joining their locations in the current proposed morphological microstate and their corresponding positions in previous accepted morphological microstate – this corrected morphological state was then used as the configuration on which to check the probability of acceptance. Typical Markov chains comprised in excess of $\mathcal{L} = 2 \times 10^6$ samples.

#### 1.3 Material parameters for hMSCs

All simulations are reported at a reference thermodynamic temperature $T = T_0$, where $T_0 = 310$ K. Most of the parameters of the model are related to the properties of the proteins that constitute stress-fibres. These parameters are thus expected to be independent of cell type. Notable exceptions to this are: (i) the stress-fibre protein volume fraction $\mathcal{H}_0$; and (ii) the passive elastic properties. Here, we use parameters calibrated for hMSCs that give good correspondence with the wide range of measurements reported in the study. The passive elastic parameters of the cytoplasm are taken to be $\mu_\mathrm{C} = 0.3$ kPa, $\kappa_\mathrm{C} = 50$ kPa and $m_\mathrm{C} = 6$, while the corresponding values for the nucleus are $\mu_\mathrm{N} = 1$ kPa, $\kappa_\mathrm{N} = 50$ kPa and $m_\mathrm{N} = 10$. The maximum contractile stress $\sigma_{\max} = 240$ kPa is consistent with a wide range of measurements on muscle fibres [6], and the density of stress-fibre proteins was taken as $\rho_0 = 3 \times 10^6$ µm$^{-3}$ with the volume fraction of stress-fibre proteins $\mathcal{H}_0 = 0.032$. Following Vigliotti et al. [2], we assume that the steady-state functional unit strain $\tilde{\varepsilon}_\mathrm{nom}^\mathrm{ss} = 0.35$ with $\mu_\mathrm{b0} - \mu_\mathrm{u} = 2.3\, k_\mathrm{B} T_0$ and $\Omega = 10^{-7.1}$ µm$^3$. The maximum angular concentration of stress-fibre proteins is set to $\hat{\eta}_{\max} = 0.8$. The cell in its undeformed state is a circle of radius $R_0$ and thickness $b_0$, with a circular nucleus of radius $R_\mathrm{N}$ whose centre coincides with that of the cell. The radius of the hMSCs in their undeformed state was taken to be $R_0 = 15$ µm, while their thickness was set at $b_0/R_0 = 0.12$. The radius of the nucleus of hMSCs in their undeformed state was taken to be $R_\mathrm{N} = 6.75$ µm, so that the nucleus occupies a volume fraction $\bar{v}_\mathrm{N} = 0.21$ of the cell.

#### 1.4 Definitions of normalised quantities and observables

Following [1], the free-energy $G^{(j)}$ can be decomposed as $G^{(j)} = \Upsilon^{(j)} + \Upsilon_0$, where $\Upsilon_0 \equiv \rho_0 V_0 \left[\mu_\mathrm{u}/n^\mathrm{R} - k_\mathrm{B} T \ln\left(\pi\, \hat{N}_\mathrm{L}\right)\right]$ is independent of the morphological microstate. It is thus natural to subtract out $\Upsilon_0$ and define a normalised free-energy as

$$\hat{G}^{(j)} \equiv \frac{\Upsilon^{(j)}}{|G_\mathrm{S} - \Upsilon_0|} = \frac{G^{(j)} - \Upsilon_0}{|G_\mathrm{S} - \Upsilon_0|}\,, \tag{1.14}$$

where $G_\mathrm{S}$ is the equilibrium free-energy of a free-standing cell (i.e. a cell in suspension with traction-free surfaces). Then, the distribution given by Eq. (1) in the main text can be re-written as

$$P_\mathrm{eq}^{(j)} = \frac{1}{\hat{Z}} \exp[-\hat{\zeta} \hat{G}^{(j)}], \tag{1.15}$$

with $\hat{Z} \equiv \sum_j \exp\left[-\hat{\zeta} \hat{G}^{(j)}\right]$ and $\hat{\zeta} \equiv \zeta |G_S - \Upsilon_0|$. It then immediately follows that the distributions of states are not influenced by the values of $n^\mathrm{R}$, $\hat{N}_\mathrm{L}$ and $V_0$ and these parameters need not be specified so long as energies are quoted in terms of the normalised energies $\hat{G}^{(j)}$. We note in passing that we have normalised the Gibbs free-energy of each morphological microstate and the homeostatic temperature by the homeostatic value of the Gibbs free-energy that the cell attains over all the morphological microstates it samples. For the hMSC parameters listed in Section 1.3, the cell in suspension has a radius of $\approx 15$ µm with a Gibbs free-energy $G_\mathrm{S} - \Upsilon_0 \approx -6.1 \times 10^9 k_\mathrm{B} T_0 = -0.03$ nJ (see Shishvan et al. [1] for details of the calculation of $G_\mathrm{S} - \Upsilon_0$). This implies that in units of Kelvin, a typical normalised temperature $1/\hat{\zeta} = 0.2$

of hMSCs corresponds to $1/\zeta \approx 10^{11}$ K. Of course, this high temperature stems from the homeostatic framework inherently recognising that the morphological fluctuations of cells, rather than having a thermal origin, are biochemical in nature and arise from the imprecise regulation of the exchange of nutrients with the nutrient bath.

Analogously, we define the normalised passive and cytoskeletal free-energies of the cell as

$$\hat{F}^{(j)}_{\text{passive}} \equiv \frac{F^{(j)}_{\text{passive}}}{|G_S - \Upsilon_0|}, \tag{1.16}$$

and

$$\hat{F}^{(j)}_{\text{cyto}} \equiv \frac{F^{(j)}_{\text{cyto}} - \Upsilon_0}{|G_S - \Upsilon_0|}, \tag{1.17}$$

respectively. Similarly, we define the normalised substrate free-energy as

$$\hat{F}^{(j)}_{\text{sub}} \equiv \frac{F^{(j)}_{\text{sub}} - \Upsilon_0}{|G_S - \Upsilon_0|}, \tag{1.18}$$

Probability density functions showing predictions of the distributions of $\hat{F}_{\text{cyto}}$ for hMSCs on selected stiffness of elastic substrates and selected areas of adhesive islands are included in Figs. 6a and 6c (in the main text), respectively. These distributions are generated by plotting histograms from the sample list generated via the MCMC procedure and normalising the frequencies to give $p(\mathrm{x})$ such that $\int_{-\infty}^{\infty} p(\mathrm{x}) d\mathrm{x} = 1$, where the dummy symbol x denotes either $\hat{F}_{\text{cyto}}$ or a morphometric variable (such as cell area, aspect ratio, and total tractions).

#### *1.4.1 Tractions exerted by cells on substrates*

An observable commonly reported in experiments, typically measured via traction-force microscopy, are spatial distributions of tractions exerted by cells on substrates and the associated so-called total traction force [7-9]. These measurements are nearly exclusively reported on relatively soft substrates where tractions exerted by the cell induce significant deformations. This makes measurements of surface displacements feasible and allows for a relatively low error estimate of the associated surface tractions [8, 9]. It is well-established that the response of cells is mechano-sensitive in the sense that the tractions they exert depend on the substrate stiffness, with the tractions increasing with increasing substrate stiffness [9, 10]. Mechanical equilibrium (Section 1.1.1) of a given morphological microstate $(j)$ specifies the traction distribution $\mathrm{T}^{(j)}(x_i)$ between the cell and the substrate. We define a normalised resultant traction

$$\hat{\mathrm{T}}^{(j)}(x_i) \equiv \frac{R_0\sqrt{\mathrm{T}_1^2 + \mathrm{T}_2^2}}{b_0 \mu_C}, \tag{1.19}$$

where $R_0$ and $b_0$ are the radius and thickness of the cell in its undeformed state, respectively, $\mu_C$ is the shear modulus of the cytoplasm and $T_i$ is the traction at location $x_i$ in morphological state $(j)$. These distributions for selected morphological microstates on elastic substrates of two stiffness ($E_{\mathrm{sub}} = 3$ kPa and 30 kPa) are included in Fig. 3d (in the main text). In addition, we also define the normalised total traction as

$$\hat{\mathrm{T}}_{\mathrm{T}}^{(j)} \equiv \frac{1}{A^{(j)}} \int_{A^{(j)}} \hat{\mathrm{T}}^{(j)} dA, \tag{1.20}$$

and the distributions of the normalised total traction are shown in Fig. 3c (in the main text) for cells on elastic substrates of three different stiffness, in Fig. 4b (in the main text) for cells on adhesive islands of three areas, and in Fig. 9c (in the main text) comparing cells untreated and treated with ROCK inhibitor.

#### *1.4.2 Construction of immunofluorescence-like images*

The predictions in Fig. 2a (in the main text) show immunofluorescence-like images comprising of three layers; the nucleus is coloured in purple and blue (in the top and bottom panels, respectively), while the focal adhesions and actin stress-fibres in shades of green and red, respectively. Focal adhesion distributions in the experiments by Engler et al [11] were observed by staining for paxillin (Fig. 2b of the main text). In the current set of simulations, the focal adhesions were not explicitly modelled but rather the cell was assumed to perfectly adhere to the substrate without directly accounting for the distribution of adhesion proteins. This results in the simulations predicting traction distributions $\mathrm{T}_i(x_i)$ as discussed above. However, it is well-known that traction magnitudes scale with concentration of adhesive proteins [12] and thus, here we use predictions of traction magnitudes as a surrogate to visualise the predictions of focal adhesion distributions. Specifically, we include in Fig. 2a (in the main text) distributions of $\hat{\mathrm{T}}^{(j)}$ for selected morphological microstates $(j)$ with the darker shades of green representing higher values of $\hat{\mathrm{T}}^{(j)}$ and thereby higher paxillin concentrations.

The predictions of the actin stress-fibre structure within the cytoplasm are coloured in red. These images, are meant to give two key pieces of information: (i) the local orientation of the dominant stress-fibre bundle, and (ii) the concentration of the stress-fibre proteins. However, no discrete fibres are present in the model of the acto-myosin stress-fibres employed here with the fibres represented via continuum internal state variables. Here, we use this continuum information to generate a discrete depiction of the fibres using the following prescription. Recall that for the purposes of calculation of $G^{(j)}$, the cell was discretized into 800 constant strain triangles of approximately equal size in the undeformed configuration. Within each element $k$, we defined an actin stress-fibre concentration in direction $\varphi^*$ as $\hat{\vartheta}_{\mathrm{b}}^{k}(\varphi^*) \equiv \hat{\eta}(\varphi^*)\hat{n}^{\mathrm{ss}}(\varphi^*)$, where $\varphi^*$ is the angle of the stress-fibre measured with respect to $x_2$ direction and is related to its angle $\varphi$ in the undeformed configuration by the rotation of element $k$. Next, we construct a dummy mesh comprising approximately 200 triangular elements to discretize the deformed cell. Elements of the original computational mesh are assigned to the element of the dummy mesh in which their centroid lies. In this manner, approximately four elements of the original computational mesh are assigned to each element of the dummy mesh.

We then define an average actin stress-fibre concentration for each element $m$ of the dummy mesh as

$$\langle \hat{\vartheta}_{\mathrm{b}}(\varphi^*) \rangle_m \equiv \frac{1}{V_m} \sum_{k \in m} \hat{\vartheta}_{\mathrm{b}}^k(\varphi^*) v_k \, , \tag{1.21}$$

where $v_k$ is the volume of element $k$ of the computational mesh that is assigned to element $m$ of the dummy mesh and $V_m = \sum_{k \in m} v_k$. The dominant stress-fibre direction $\varphi^*_{\mathrm{max}}$ in dummy element $m$ is defined as the value of $\varphi^*$ corresponding to the maximum value of $\langle \hat{\vartheta}_{\mathrm{b}}(\varphi^*) \rangle_m$. Each element of the dummy mesh is hatched with green lines in the direction $\varphi^*_{\mathrm{max}}$. We set the spacing of the hatches to scale with the local density of bound stress-fibre proteins in order to replicate the higher intensity of the fluorescent phalloidin observed under light microscopy in locations with a higher concentration of stress-fibres. The total concentration of bound stress-fibre proteins in element $k$ of the computational mesh is

$$\hat{N}_{\mathrm{b}}^k = \int_{-\pi/2}^{\pi/2} \hat{\vartheta}_{\mathrm{b}}^k(\varphi) \, d\varphi \, , \tag{1.22}$$

with $\hat{N}_{\mathrm{b}}^{\mathrm{T}} \equiv \int_{V_{\mathrm{C}}} \hat{N}_{\mathrm{b}} \; dV$ denoting the total concentration of bound stress-fibre proteins in the cell.

The spacing $\mathcal{s}$ of the green hatches in element $m$ of the dummy mesh is set to scale inversely with the concentration of bound stress-fibre proteins within that element, i.e. $\mathcal{s} \equiv \hat{N}_{\mathrm{b}}^T / \sum_{k \in m} \hat{N}_{\mathrm{b}}^k$ . This process results in the predictions of the actin cytoskeletal structure shown in Fig. 2a (in the main text), and we include in the top panel of Fig. 10b (in the main text) only predictions of the actin cytoskeletal structure (i.e. without nucleus or focal adhesions) for cells on adhesive islands of area $A_{\mathrm{p}} = 2725 \, \mu m^2$ to give more insight into the details of the predicted cytoskeletal structure at various locations within the cell. The predictions in Fig. 4a (in the main text) are generated by the same method, with the nucleus coloured in blue, while the focal adhesions (i.e. distributions of $\hat{\mathrm{T}}^{(j)}$) and actin stress-fibres are coloured in shades of pink and green, respectively.

### References


1. Shishvan SS, Vigliotti A, Deshpande VS. 2018 The homeostatic ensemble for cells. *Biomech Model Mechanobiol,* **17(6)**:1631-1662.
2. Vigliotti A, Ronan W, Baaijens FPT, Deshpande VS. 2015 A thermodynamically motivated model for stress-fibre reorganization. *Biomech. model. Mechanobiol.* **15**:761-789.
3. Buskermolen ABC, Suresh H, Shishvan SS, Vigliotti A, DeSimone A, Kurniawan NA, Bouten CVC, Deshpande VS. 2019 Entropic forces drive cellular contact guidance. *Biophys. J.,* **116**, 1994-2008.
4. Ogden RW. 1972 Large Deformation Isotropic Elasticity – On the correlation of theory and experiment for incompressible rubberlike solids, *Proc. Royal Soc. London A, Math. Phys. Sci.* **326**:565-584.
5. Metropolis N, Rosenbluth AW, Rosenbluth MN, Teller AH, Teller E. 1953 Equation of state calculations by fast computing machines. *J. Chem. Phys.* **21**:1087-1092.
6. Lucas SM, Ruff RL, Binder MD. 1995 Specific tension measurements in single soleus and medial gastrocnemius muscle fibres of the cat. *Exp. Neurology* **95**:142-154.

7. Tan JL, Tien J, Pirone DM, Gray DS, Bhadriraju K, Chen CS. 2003 Cells lying on a bed of microneedles: An approach to isolate mechanical force. *Proc. Natl. Acad. Sci. USA.* **100**:1484-1489.
8. Maskarinec SA, Franck C, Tirrell DA, Ravichandran G. 2009 Quantifying cellular traction forces in three dimensions. *Proc. Natl. Acad. Sci. USA.* **106**:22108-22113.
9. Califano JP, Reinhart-King CA. 2010 Substrate stiffness and cell area predict cellular traction stresses in single cells and cells in contact. *Cell. Mol. Bioeng.* **3(1)**:68–75.
10. Oakes PW, Banerjee S, Marchetti MC, Gardel ML. 2014 Geometry regulates traction stresses in adherent cells. *Biophys J.* **107(4)**:825–833.
11. Engler AJ, Sen S, Sweeney HL, Discher DE. 2006 Matrix elasticity directs stem cell lineage specification. *Cell* **126**, 677–689.
12. Dumbauld DW, Lee TT, Singh A, Scrimgeour J, Gersbach CA, Zamir EA, Fu J, Chen CS, Curtis JE, Craig SW, García AJ. 2013 How vinculin regulates force transmission. *Proc. Natl. Acad. Sci. USA.* **110**:9788-9793.
